# Aquaporin-4 Regulates Edema, Sustains Astrocyte Differentiation, and Constrains Immunosuppressive Remodeling in Glioblastoma

**DOI:** 10.64898/2026.10.23.761345

**Authors:** Tanvi Joshi, Kaisha Garvin, Matthew J. Flick, Montserrat Puigdelloses Vallcorba, Mehek Dedhia, Zhihong Chen, Raina Roche, Emir Radkevich, Susana Isabel Ramos, Nadejda M. Tsankova, Nhan L. Tran, Leland S. Hu, Katherine B. Peters, Alex J. Smith, Dieter Henrik Heiland, Dolores Hambardzumyan

**Affiliations:** Department of Oncological Sciences, The Tisch Cancer Institute, Icahn School of Medicine at Mount Sinai, New York NY, USA; Department of Cancer Biology, Mayo Clinic, Phoenix AZ, USA; Department of Radiology, Mayo Clinic, Phoenix AZ, USA; Icahn School of Medicine at Mount Sinai, New York NY, USA; Human Immune Monitoring Center, Icahn School of Medicine at Mount Sinai, New York NY, USA; Department of Neurosurgery, Preston Robert Tisch Brain Tumor Center, Duke University School of Medicine, Durham NC, USA; Department of Neuroscience and Department of Pathology, Molecular and Cell Based Medicine, Icahn School of Medicine at Mount Sinai, New York NY, USA; Department of Neurological Surgery, Mayo Clinic, Phoenix AZ, USA; Department of Ophthalmology, University of California, San Francisco, San Francisco CA, USA; Translational Neurosurgery, Friedrich-Alexander-Universität Erlangen-Nürnberg, Erlangen, Germany; Department of Neurosurgery, University Hospital Erlangen, Friedrich-Alexander-Universität Erlangen-Nürnberg, Erlangen, Germany; Department of Neurological Surgery, Northwestern University Feinberg School of Medicine; Department of Neurosurgery, Icahn School of Medicine at Mount Sinai, New York NY, USA; Department of Neurosurgery, Perelman School of Medicine, University of Pennsylvania, Philadelphia PA, USA

## Abstract

Vasogenic cerebral edema is a severe complication that affects most patients with glioblastoma (GBM). It leads to an increase in intracranial pressure, neurological deficits, and increased mortality rates. While there is considerable research on edema formation, the molecular and cellular mechanisms of edema resolution in GBM, particularly in response to dexamethasone or other anti-edema therapies, remain largely undefined. Our findings demonstrate that the water channel aquaporin-4 (AQP4), a key regulator of brain water homeostasis, is upregulated in human GBM compared to the normal brain. AQP4 expression correlates with peritumoral edema in GBM patients, with the classical (CL) GBM subtype exhibiting the highest *AQP4* levels but the lowest edema content relative to the proneural (PN) and mesenchymal (MES) subtypes. Using the RCAS/*tv-a* system, we generated *de novo* GBM with defined driver mutations in *Aqp4*-proficient and -deficient mice. Loss of *Aqp4* shortened survival and exacerbated edema, accompanied by reduced astrocytic coverage within the tumor core, but not at the tumor margin. Single-cell RNA-sequencing and spatial transcriptomics revealed that *Aqp4* loss reprograms astrocytes toward an immature, progenitor-like state and alters their crosstalk with other cell types in the GBM microenvironment. Additionally, loss of *Aqp4* skewed myeloid populations toward immunosuppressive phenotypes while reducing lymphoid infiltration. Treatment with anti-VEGFA antibody, an established anti-edema therapy, increased expression of *Aqp4* and reduced vessel leakage, underscoring the role of AQP4 in the therapeutic modulation of brain water balance. Overall, this study highlights the important role of AQP4 in resolving edema and its critical function in maintaining astrocyte identity and shaping the immune landscape of GBM.

## Introduction

Glioblastoma (GBM) is the most common and aggressive primary brain tumor in adults with median survival ranging from 13-24 months and a dismal 5-year survival rate of ∼6.8% despite standard of care treatment involving surgical resection, radiation and chemotherapy with temozolomide (TMZ) (1–3). One of the major complications observed in GBM patients is cerebral edema, defined as a pathological increase in fluid accumulation in the brain, often exceeding tumor mass. Due to the physical limitations imposed by the rigid skull, cerebral edema and tumor mass both contribute to increased intracranial pressure and accompanying neurological deficits (4). Several retrospective studies have shown that the extent of contrast enhancement on a T2-weighted MRI scan, indicative of peritumoral brain edema, is a prognostic indicator of poor overall survival in GBM patients and severely impacts their quality of life (5, 6). End-stage GBM-associated edema is typically vasogenic in nature, resulting from enhanced vascular permeability due to a compromised blood-brain barrier (BBB). While the mechanism by which GBM-associated edema accumulates is more known, edema resolution in GBM is not well understood. Current management strategies for GBM-associated edema rely primarily on the corticosteroid dexamethasone (DEX) and, in some cases, anti-VEGF therapies (7–12). While these agents reduce edema, their mechanisms of action remain poorly understood, and the drawbacks associated with their use highlight the need to both reconsider their utilization and advance our mechanistic understanding of anti-edema therapies in GBM, especially as it pertains to water resolution (13–20).

The human brain consists of approximately 80% water (21). The key regulators of water movement and homeostasis in the brain are the aquaporins, with aquaporin-4 (AQP4) being the predominant water channel in the brain. AQP4 at the astrocytic endfeet regulates bidirectional movement of water across the blood-brain and brain-CSF interfaces (22). Early studies on AQP4 in pathological conditions uncovered a dual role for the water channel in the formation of cytotoxic edema, where the BBB is intact, and in the resolution of vasogenic edema, where the BBB is compromised (23, 24). The models used to study AQP4 in vasogenic edema involved intraparenchymal fluid infusion, cortical freeze injury, and metastatic tumor cell implantation in the brain. Studies on the role of AQP4 in gliomas are very limited and largely correlative, showing increased expression of AQP4 in human GBM samples compared to the normal brain and low-grade gliomas (25–27), or they are based on *in vitro* experiments which do not capture the dynamic tumor microenvironment and complexity of edema formation (28, 29).

Astrocytes are the primary cell type expressing AQP4 in the brain and are the most abundant non-immune cells in the GBM microenvironment (30). The advent of improved isolation techniques for astrocytes along with single-cell sequencing and spatial transcriptomics technologies have allowed high-resolution investigations into the roles of these glial cells in health and disease, revealing them to be active participants in tumor progression rather than passive bystanders. In addition to paracrine signaling, astrocytes directly exchange metabolites and molecular cargo with tumor cells through gap junctions and extracellular vesicles, further augmenting tumor fitness (31–33). They also remodel the extracellular matrix and perivascular niche, facilitating tumor cell infiltration and modulating vascular permeability (34, 35). However, the roles of AQP4 in regulating astrocyte responses, and their interactions with other cell types in the GBM microenvironment, remain to be studied.

In this study, we investigated how AQP4 influences edema formation and immune modulation in GBM using a combination of human GBM tissue and genetically engineered mouse models (GEMM) of *de novo* GBM in *Aqp4*-proficient and -deficient backgrounds. Analysis of human GBM samples revealed a negative correlation between *AQP4* expression and MRI-based edema burden. In murine models, *Aqp4* loss decreased survival and exacerbated peritumoral edema, suggesting that AQP4 facilitates edema resolution. Mechanistically, AQP4 expression localized primarily to tumor-associated astrocytes rather than tumor cells, and its loss reduced astrocytic coverage of the tumor core and promoted an immature, progenitor-like astrocyte phenotype. Integrative single-cell RNA sequencing (scRNA-seq), spatial transcriptomics and flow cytometry revealed that *Aqp4* deficiency reshapes the immune microenvironment, by depleting lymphoid populations, enriching immunosuppressive tumor-associated macrophages (TAM), and inducing neuronal and synaptic gene programs at the tumor margin. Importantly, *Aqp4* loss enhanced vascular permeability, which was mitigated by anti-VEGF therapy in an *Aqp4*-dependent manner. Together, these findings identify AQP4 as a key regulator of edema and vascular integrity in GBM, while also uncovering its previously unrecognized roles in astrocyte function and immune microenvironment remodeling, highlighting AQP4 as a potential therapeutic node at the intersection of tumor biology, immunity, and brain homeostasis.

## Results

### *AQP4* expression negatively correlates with edema content in human GBM

Peritumoral edema detected on T2-weighted MRI is a well-established independent predictor of poor overall survival in patients with GBM (5, 6). However, the relationship between *AQP4* expression and GBM-associated edema has not been clearly defined. To investigate this, we analyzed previously published bulk RNA sequencing data from multiregional GBM samples (n=159) with spatially matched regional measurements from contrast-enhanced MRI (36). Each biopsy was precisely localized to its corresponding position on the patient’s preoperative T2-weighted MRI scan to determine its regional signal intensity. Biopsies sampled from regions T2 hyperintense (Fig. 1A, red) regions reflected areas of higher water content, whereas isointense regions (Fig. 1A, blue) represented lower water content. This strategy enabled direct correlation of *AQP4* expression with MRI signal intensity within the same tumor.

**Fig. 1.**
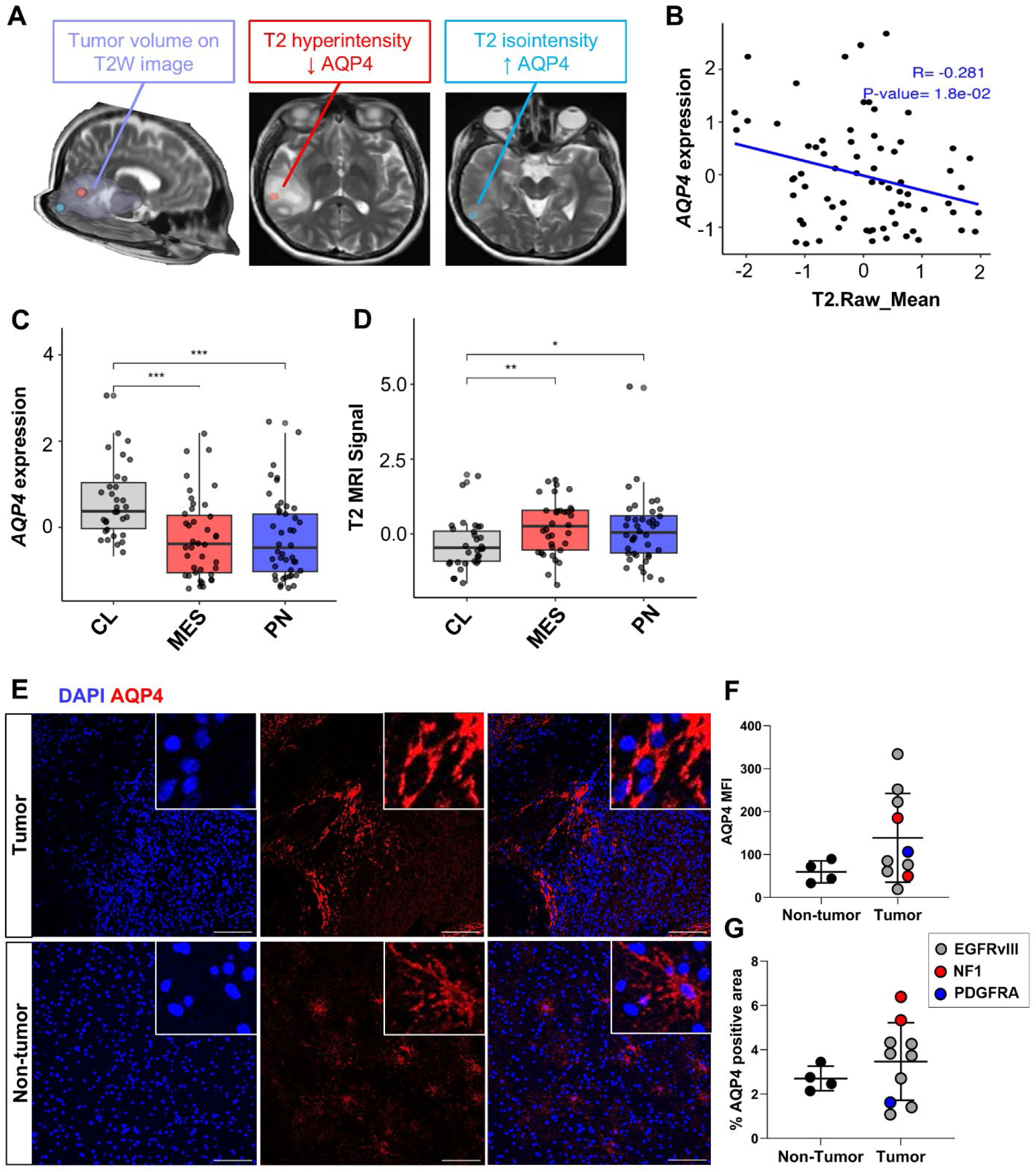
AQP4 expression negatively correlates with MRI signal in human GBM. **A**, Representative tumor volume on T2-weighted (T2W) image showing two multi-regional intraoperative biopsy sites from a tumor mass lesion (left, purple); biopsy of a T2 hyperintense tumor region with low *AQP4* expression (middle, red); separate biopsy of a T2 iso-intense region with high *AQP4* expression (right, blue). **B**, Scatter plot depicting the association between z-score normalized *AQP4* expression and T2 signal intensity across all samples. Each point represents an individual sample. The blue line represents the linear regression fit. Pearson’s correlation analysis (R = –0.281, P = 0.018). **C&D**, Box-and-whisker plots showing the distribution of *AQP4* expression and T2-weighted MRI intensity across Verhaak subtypes (CL-Classical, MES - Mesenchymal, PN - Proneural). Each point represents an individual sample. Global significance was tested using Type II ANOVA, and pairwise post-hoc comparisons were performed using the *emmeans* R package. *p < 0.05, **p < 0.01, ***p < 0.001. **E**, Representative images from immunofluorescence staining for AQP4 (red) in human GBM and non-tumor tissue sections. Images were taken on a Leica SP8 confocal microscope. Scale bar=100 μm. **F**, Quantification of AQP4 mean fluorescent intensity (MFI) in non-tumor (n=4) and tumor tissue sections (n=6). Values were compared using two-tailed Student’s t-test. **G,** Quantification of AQP4 positive area in non-tumor (n=4) and tumor (n=6) tissue sections. Values were compared using two-tailed Student’s t-test.

Pearson correlation analysis revealed a significant negative correlation between z-score-normalized *AQP4* expression and T2-weighted MRI intensity (R = –0.281, p = 0.018) among contrast-enhancing samples (Fig. 1B). These findings complement previous reports showing an inverse correlation between AQP4 and proton density (PD) in peritumoral regions of patients with meningioma, GBM, and brain metastases (37). Given that GBM transcriptional subtypes exhibit distinct molecular and histopathologic features, we next investigated the relationship between *AQP4* expression and MRI signal intensity by Verhaak subtype (38, 39). The Classical (CL) subtype displayed the highest *AQP4* expression and lowest T2 signal intensity, while the Mesenchymal (MES) and Proneural (PN) tumors showed lower *AQP4* expression and correspondingly higher edema burden (Fig. 1C&D). Because dexamethasone (DEX) is routinely administered to manage cerebral edema in GBM, we further examined whether steroid treatment influences AQP4 protein expression. In a separate patient cohort, AQP4 mean fluorescent intensity (MFI) and positive area were quantified in non-tumor and tumor tissue (Fig. 1E–G). Correlative analysis showed that higher AQP4 MFI was associated with lower daily DEX dose, whereas AQP4-positive area exhibited no significant relationship with DEX dose (Supp. Fig. 1A-C). Notably, previous studies have reported that DEX can variably modulate AQP4 expression depending on tissue context and inflammation (40–48). Collectively, these results indicate that *AQP4* expression inversely correlates with edema burden in human GBM, supporting a role for AQP4 in edema resolution and suggesting that DEX treatment may influence AQP regulation, an important consideration when interpreting AQP4 levels in clinical specimens.

### Loss of *Aqp4* results in decreased survival, increased edema content, and reduced tumor volume in GBM-bearing mice

Based upon our observations in human GBM patient data, which showed a negative correlation between *AQP4* expression and T2 MRI signal, we set out to further investigate the role of this water channel in tumor-bearing mice (Fig. 2A&B). To generate *de novo* GBM in mice, we employed RCAS/*tv-a* (Replication-Competent Avian Sarcoma-Leukosis virus/tumor virus receptor-A), a somatic cell type-specific gene transfer system that allows for introduction of combinations of driver mutations commonly seen in human GBM, along with silencing of tumor suppressor genes into a target cell population (49, 50). First, *Aqp4^-/-^* mice were crossed to *Nestin tv-a* (*Ntv-a*) mice to generate *Ntv-a;Aqp4^-/-^* mice. Using a combination of RCAS-*PDGFB* (overexpression of *PDGFB*) and RCAS-shRNA*-p53*, we generated murine GBM (referred to as PDGFB mGBM) that closely resembles the human PN GBM subtype (51, 52) in *Ntv-a;Aqp4^+/+^*(*Aqp4*-proficient) and *Ntv-a;Aqp4^-/-^* (*Aqp4*-deficient) mice. A combination of *Nf1, p53,* and *Pten* silencing, along with overexpression of *PDGFA,* was used to generate mGBM that closely resembled MES human GBM (Nf1 mGBM) in *Aqp4*-proficient and *Aqp4*-deficient mice. We also generated mGBM that resembled the human CL GBM subtype (EGFRvIII mGBM) in *Aqp4*-proficient mice as previously described (51) (Supp. Fig. 2A). We first performed qPCR and immunofluorescence staining for AQP4 in *Aqp4*-proficient tumor-bearing and naïve mice. Our results showed that tumor-bearing mice had elevated levels of AQP4 at both the transcriptional and protein levels compared to naïve mice (Supp. Fig. 2B, E&F). While the AQP4 staining in normal brains exhibited a vascular pattern, tumor-bearing brains had a largely ubiquitous pattern of AQP4 expression. While we did not observe any significant differences in *Aqp4* expression across the different mGBM subtypes, we observed varying levels of edema content with PDGFB and EGFRvIII GBM-bearing mice showing higher edema content compared to Nf1 GBM-bearing mice (Supp. Fig. 2C&D). Kaplan-Meier survival analysis showed a decrease in median survival in *Aqp4*-deficient mice compared with *Aqp4*-proficient tumor-bearing mice in both PDGFB mGBM (m.s 44 days vs. 51.5 days) and Nf1 mGBM (m.s 61 days vs. 72.5 days) models (Fig. 2B & F). Next, we measured edema content in *Aqp4*-proficient and *Aqp4*-deficient tumor-bearing mice using the wet/dry assay. Because severe fluid accumulation in human GBM typically indicates advanced or terminal disease, edema content in tumor-bearing mice was measured at the onset of endpoint signs. In both mGBM models, *Aqp4*-deficient mice had higher edema content compared to wild-type (WT) mice (Fig. 2C&G). PDGFB mGBM-bearing *Aqp4*-proficient mice had higher edema content compared to Nf1 mGBM-bearing mice (Supp. Fig. 2D), potentially owing to the abnormal and highly permeable vasculature that defines the PDGFB mGBM model (51, 53, 54). To assess the effect of *Aqp4* loss on tumor volume, we also performed MRI on end-stage tumor-bearing mice using a T2 sequence on a 9.4T Bruker small animal scanner. Interestingly, in both models, we observed decreased tumor volume in *Aqp4*-deficient mice compared to WT mice (Fig. 2D, E, H & I). As we have previously shown (13, 14), end-stage tumor burden is the sum of tumor volume and edema measured on T2-weighted MRI scans; this suggests that decreased tumor volume in *Aqp4*-deficient mice results from increased edema content. We further confirmed these results by serially sectioning brains for tumor volume reconstitution at the onset of endpoint signs using an established protocol (14, 55) (Supp. Fig. 3A-C). Since DEX treatment reduces edema but does not prolong survival in tumor-bearing mice (13, 14), this raises the question of whether increased edema alone can drive shortened survival in *Aqp4*-deficient mice, or if additional factors also contribute. To determine whether there were differences in tumor proliferation that would contribute to shorter survival in *Aqp4*-deficient mice compared to WT mice, we next performed immunohistochemical (IHC) staining for phosphor-histone H3 (pH3 – a marker of cells in the M-phase of the cell cycle) in tumor sections from PDGFB and Nf1 mGBM-bearing *Aqp4*-proficient and *Aqp4*-deficient mice (53, 56). We did not observe differences in the numbers of pH3-positive cells in any of the groups (Supp. Fig. 3D&E). These results indicate that the increased edema in mice lacking AQP4 channels could be imposing physical constraints on tumor growth while also driving the survival difference observed in comparison to *Aqp4*-proficient mice. It is plausible that mechanical constraints imposed by the skull, coupled with elevated intracranial pressure, induce axonal injury, which may contribute to reduced survival, as recently demonstrated (57). Since fluctuations in fluid accumulation as well as tumor volume directly impact intracranial pressure (ICP) in the brain, we established an ICP measuring system for GBM-bearing mice. First, we measured ICP in naïve *Aqp4*-proficient and *Aqp4*-deficient mice and demonstrated higher ICP in *Aqp4*-deficient mice compared to WT mice (Fig. 2 J&K). Next, we measured ICP in tumor-bearing mice. Tumor-bearing WT mice in PDGFB mGBM had higher ICP compared to naïve mice (Fig. 2J&L). *Aqp4*-proficient and *Aqp4*-deficient mice bearing PDGFB mGBM did not differ in their mean ICP values (Fig. 2L&M). Overall, these results suggest that AQP4 is involved in edema resolution from GBM, as loss of the water channel in tumor-bearing mice results in increased edema leading to physical limits on tumor growth as well as shortened survival. We also established an ICP measurement protocol as a secondary readout for edema and tumor volume-related changes in GBM-bearing mice. While the protocol demonstrates high sensitivity in detecting subtle genotype-dependent alterations, it remains sufficiently robust to reliably distinguish tumor-bearing from naïve conditions.

**Fig. 2:**
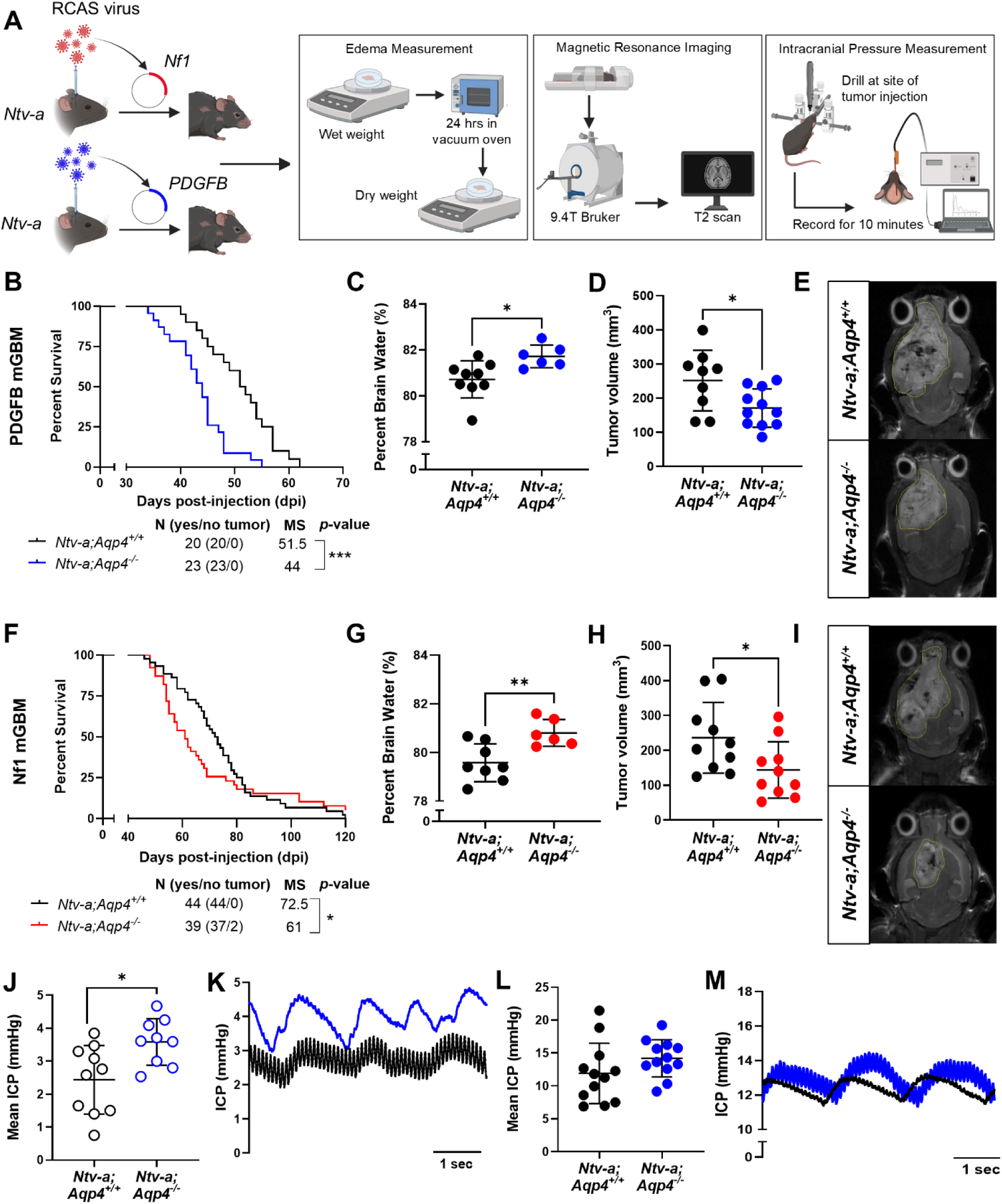
Loss of *Aqp4* results in decreased survival, increased edema content but reduced tumor burden in tumor-bearing mice. A, Schematic illustration of MRI, edema and intracranial pressure measurement in tumor-bearing mice. **B and F**, Kaplan-Meier survival curves demonstrating decreased median survival of *Ntv-a;Aqp4^-/-^* mice bearing PDGFB and Nf1 mGBM tumors, *p<0.05; ***p<0.001 by log-rank (Mantel-Cox) test. **C and G**, Graphs comparing edema content as percent brain water in *Ntv-a;Aqp4^+/+^* and *Ntv-a;Aqp4^-/-^* mice bearing PDGFB and Nf1 mGBM tumors. Two-tailed Student’s t-test, *p<0.05; **p<0.01. **D and H**, Comparison of endpoint tumor burden between *Ntv-a;Aqp4^+/+^* and *Ntv-a;Aqp4^-/-^* mice bearing PDGFB and Nf1 mGBM tumors by T2-weighted MRI. Two-tailed Student’s t-test, *p<0.05. **E and I**, Representative T2-weighted MRI scans indicating a decrease in endpoint tumor burden in *Ntv-a;Aqp4^-/-^* tumor-bearing mice as compared to *Ntv-a;Aqp4^+/+^* mice. **J and K**, Comparison of intracranial pressure (ICP) between naïve *Ntv-a;Aqp4^+/+^* and *Ntv-a;Aqp4^-/-^* mice and representative ICP traces showing cardiac (peak) and respiratory (trough) pulsations. Two-tailed Student’s t-test, *p<0.05. **L and M**, Comparison of intracranial pressure (ICP) between *Ntv-a;Aqp4^+/+^* and *Ntv-a;Aqp4^-/-^* mice bearing PDGFB mGBM and representative ICP traces showing cardiac (peak) and respiratory (trough) pulsations.

### AQP4 is primarily expressed by astrocytes in the GBM microenvironment

Early studies that were focused on AQP4 in the context of human glioma tissue employed IHC staining to assess the expression levels of the water channel. These studies were limited to only showing that AQP4 expression is higher and more ubiquitous in glioma tissue as compared to the normal brain, and that it increases with tumor grade (25–27). While astrocytes are known to express AQP4 on their perivascular endfeet (58), GBM cells also display phenotypic plasticity, often adopting markers that are typically associated with other cell types in the microenvironment (59). Additionally, some *in vitro* studies have reported AQP4 expression on glioma cell lines (28, 29). To determine the source(s) of AQP4 in murine and human GBM, we utilized a combination of RNAScope and multiplex immunofluorescent (IF) staining on tumor tissue sections. In the RNAScope experiment, AQP4-positive areas overlapped with GFAP-positive reactive astrocytes, but not with OLIG2-positive tumor cells (Fig. 3A&B). We confirmed this observation at the protein level in PDGFB mGBM tissue through co-localization analysis with a combination of anti-GFAP and anti-AQP4 antibody staining (Fig. 3C-E). We also performed scRNA-seq on PDGFB mGBM tissue from *Aqp4*-proficient mice (n=4) and found that *Aqp4* expression coincided with the expression of classic astrocyte markers such as *Gfap* as well as *Aldh1l1* (Supp. Fig. 4A-C). To further validate these findings in mGBM, we performed western blot analysis on a combination of mGBM-derived tumor tissue and primary cultures (Fig. 3I). While all tumor-derived snap-frozen tissue samples demonstrated AQP4 expression, none of the tumor-derived primary cultures showed AQP4 positivity (Fig. 3J). AQP4 positivity was also present in primary astrocyte cultures derived from postnatal day 0 (P0)-P3 pups. Since tumor-derived primary cultures are exclusively representative of tumor cells and the tumor-derived tissue includes other cell types in the GBM microenvironment such as AQP4-expressing astrocytes, these results indicate that AQP4 is primarily expressed on astrocytes and not by tumor cells. To investigate whether these findings hold true in the setting of human GBM, we performed multiplex immunofluorescent staining using the Orion^™^ platform on human GBM samples (n=6) with *EGFRvIII* mutation, *NF1* mutation or *PDGFRA* amplification. By quantifying co-localization of AQP4 with GFAP-positive and OLIG2-positive cells from the Orion^™^ panel, we demonstrated that GFAP-positive cells show the highest level of co-localization with AQP4 (Fig. 3F-H and Extended Data Fig. 1A-C). In contrast to murine GBM, we additionally observed low levels of AQP4-positivity in other cell types, including OLIG2-positive cells, tumor-associated macrophages (TAMs) and T cells (Extended Data Fig. 1C-F). Similar to our mouse models, we also performed western blot analysis of human non-tumor cortex, GBM-derived tumor tissue and primary cultures (Fig. 3K). Our observations mirrored our findings in mGBM, where tumor-derived tissue samples expressed AQP4 while tumor-derived primary cultures did not, suggesting that tumor cells are not the main source of AQP4 (Fig. 3L).

**Fig. 3:**
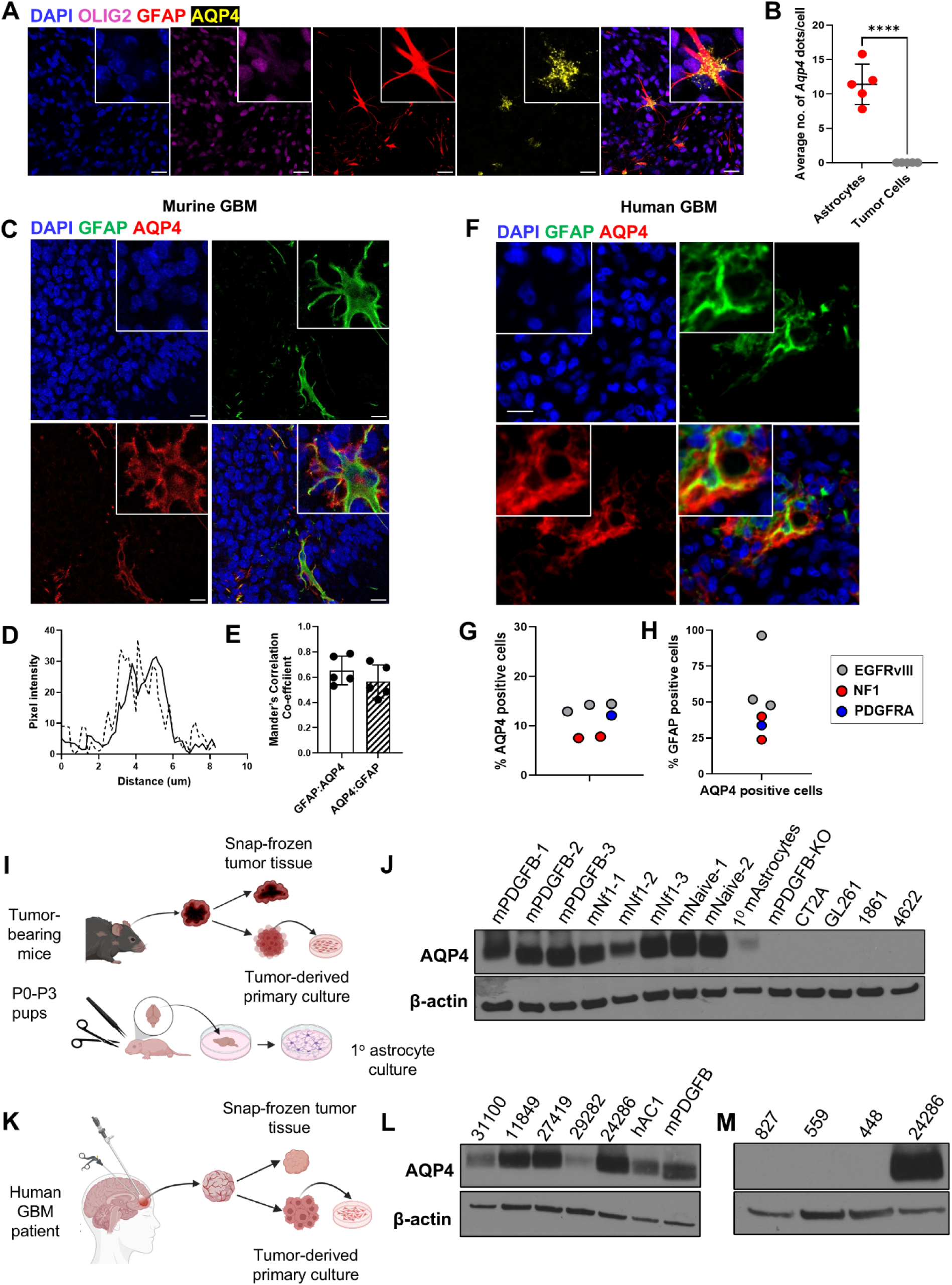
AQP4 co-localizes with GFAP-positive reactive astrocytes in murine GBM. **A**, Representative images from RNAscope in-situ hybridization for mRNA of *Aqp4* (yellow), combined with immunofluorescence staining for OLIG2-positive tumor cells (purple) and GFAP-positive reactive astrocytes (red) in PDGFB mGBM tissue. DAPI was used as a nuclear stain. Images were taken on the Leica Stellaris 5 confocal microscope. Scale bar=20 μm. **B**, Quantification of the number of *Aqp4* mRNA dots co-localizing with OLIG2-positive tumor cells and GFAP-positive reactive astrocytes. At least 3 images were quantified per sample. Two-tailed Student’s t-test ****p<0.0001. **C**, Representative images of immunofluorescence staining for GFAP (green) and AQP4 (red) in PDGFB mGBM samples. DAPI was used as a nuclear stain. Images were taken on a Leica SP8 confocal microscope. Scale bar=20 μm. **D and E**, Co-localization analysis plot and graph showing Mander’s correlation coefficient of GFAP and AQP4 across an ROI using the Coloc 2 function in ImageJ with pixel intensity on the Y-axis and distance in μm on X-axis. Several ROIs were outlined per sample and n=5 samples were used for analysis. **F**, Representative images of immunofluorescence staining for GFAP (green) and AQP4 (red) in human GBM tissue samples using the Orion platform. DAPI was used as a nuclear stain. Scale bar=20 μm. **G and H**, Quantification of AQP4 positive cells (left) and AQP4 positive cells as a percentage of GFAP positive cells (right) in human GBM tissue sections (n=6). **I and K**, Schematic illustration indicating isolation of murine and human GBM tissue and tumor-derived primary cultures respectively for Western blot-based detection of AQP4 protein levels. **J**, Western blot for AQP4 for, from left to right, murine PDGFB-driven GBM tissue (mPDGFB1-3), shNf1-driven GBM tissue (mNf1-1-3), naïve brain tissue (mNaive1-2), primary astrocyte culture (1° mAstrocytes), PDGFB mGBM tissue from *Ntv-a;Aqp4^-/-^* mice (negative control) and tumor-derived primary cultures (mGSC1-4). Beta-actin was used as a loading control. **L**, Western blot for AQP4 for human proneural GBM tissue (11849), mesenchymal GBM tissue (31100, 27419), classical GBM tissue (29282, 24286) and naïve adult cortex tissue (hAC1). PDGFB mGBM tissue (mPDGFB) was used as technical control and beta-actin was used as a loading control. **M**, Western blot for AQP4 in human GBM-derived primary cultures (827, 559, 448). Human classical GBM tissue (24286) was used as a technical control. Beta-actin was used as a loading control.

### Loss of *Aqp4* results in decreased astrocyte reactivity within the tumor core, but not at the tumor margin

While reactive astrocytes have been well-studied and are often implicated in promoting GBM progression (60–62), the mechanisms driving their activation within the tumor context remain largely understudied. Therefore, we next evaluated the effects of *Aqp4* loss on astrocyte reactivity and morphology in GBM. We performed IF staining for GFAP in PDGFB and Nf1 mGBM generated in *Aqp4-*proficient and *Aqp4-* deficient mice. The results indicate that PDGFB mGBM from *Aqp4*-deficient mice showed a drastic decrease in GFAP-positive area compared to tumors from WT mice (Fig. 4A&B). The typical reactive morphology of astrocytes, including an enlarged cell body and increased branching of processes, observed in tumors from *Aqp4*-proficient mice was lost in tumors from *Aqp4*-deficient mice. Additionally, this decrease was evident only in the tumor core, not at the tumor margin. Similar results were obtained in Nf1 mGBM-bearing mice (Supp. Fig. 5A&B). We additionally validated these results using the *Aldh1l1-eGFP/Rpl10a* mouse line crossed to *Aqp4*-proficient or *Aqp4*-deficient mice (Fig. 4C-E) (63–65). AQP4 is typically found on astrocytic endfeet in perivascular areas in the normal brain (58). To assess if loss of *Aqp4* influenced perivascular astrocyte coverage, we stained for GFAP and CD31 (blood vessel marker) in tissue sections from *Aqp4-*proficient and *Aqp4-*deficient mice bearing PDGFB mGBM (Supp. Fig. 5C). As observed in other IF experiments, there was a decrease in perivascular GFAP-positive area in the *Aqp4*-deficient mice compared to WT mice (Supp. Fig. 5D&E). While IF staining for markers such as GFAP and ALDH1L1 allowed us to study astrocyte coverage and morphology, it did not provide information about whether changes in astrocyte numbers also contribute. Thus, to investigate whether the numbers of astrocytes were affected in *Aqp4*-deficient mice, we performed flow cytometry to gate ALDH1L1-eGFP-positive astrocytes in naïve and PDGFB mGBM-bearing mice. We did not observe any significant differences in the GFP-positive astrocyte population in either group (Fig. 4F). These results indicated that while loss of *Aqp4* had a significant effect on astrocyte reactivity, it did not affect their numbers in GBM. To further study transcriptional changes in the tumor microenvironment induced by *Aqp4* loss, we performed scRNA-seq on PDGFB mGBM tumors from *Aqp4-*proficient (n=4) and *Aqp4-*deficient (n=3) mice. We generated a scRNA-seq dataset comprised of 99,233 high-quality cells, each with an average of 8466 unique molecular identifiers (UMIs) and 2554 detected genes per cell. We integrated the dataset using Harmony, followed by unsupervised clustering and automated as well as manual cell-type annotation. The final cell-type classification was achieved by merging automated labels with clustering results, revealing a clear separation of major cell types, including malignant cells (AC-, OPC-, MES-, and NPC-like states) (66), myeloid cells (microglia and bone marrow-derived myeloid cells or BMDMs), astrocytes as well as other glial cells, lymphoid cells and cell types associated with the neurovascular unit (pericytes and endothelial cells) (Fig. 4G). Next, we looked at the relative abundance of each cell type between PDGFB mGBM generated in *Aqp4-*proficient and *Aqp4-*deficient mice, demonstrating an enrichment of tumor cell clusters in *Aqp4*-deficient tumors (Fig. 4H&I). In contrast, WT tumors exhibited increased T cells and myeloid cells, suggesting a more pronounced inflammatory response. Differential gene expression analysis of the astrocyte subcluster revealed that *Aqp4-*deficient astrocytes upregulated *Apod, Mbp, Mobp*, and *Nd4l*, while downregulating immune-related genes such as *Hla-a* and *Hla-b* (Fig. 4J). Gene set enrichment analysis confirmed that astrocytes from *Aqp4-*proficient tumors exhibited enrichment of immune-related pathways, including ‘immune response,’ ‘peptide antigen processing and presentation,’ and ‘MHC protein complex’ (Supp. Fig. 6A). In contrast, *Aqp4-*deficient astrocytes were enriched for pathways related to ‘cadherin binding in cell-cell adhesion,’ ‘cell cycle checkpoint regulation,’ and metabolic processes such as ‘long-chain fatty acid metabolism’ (Supp. Fig. 6B). To further characterize the functional state of astrocytes, we applied an astrocyte subtyping framework as described by Heiland *et al.* (30). Astrocytes within the tumor microenvironment, termed tumor-associated astrocytes (TAAs), adopt specialized reactive states that extend beyond conventional gliosis. In this model, TAAs differentiate along a trajectory from a progenitor-like to a mature state (Y-axis) and from an inflammatory response to a wound-healing STAT3-activated phenotype (X-axis). We observed a shift towards a progenitor-like state in *Aqp4-*deficient astrocytes, accompanied by an enrichment of the proliferative marker, *Top2a*, which is typically upregulated in immature or fetal astrocytes (67) (Fig. 4K&L). Together, these results suggest that *Aqp4* loss leads to a decrease in astrocyte reactivity and a shift towards an immature phenotype, likely influencing the tumor microenvironment by dampening inflammation and enhancing tumor cell persistence.

**Fig. 4:**
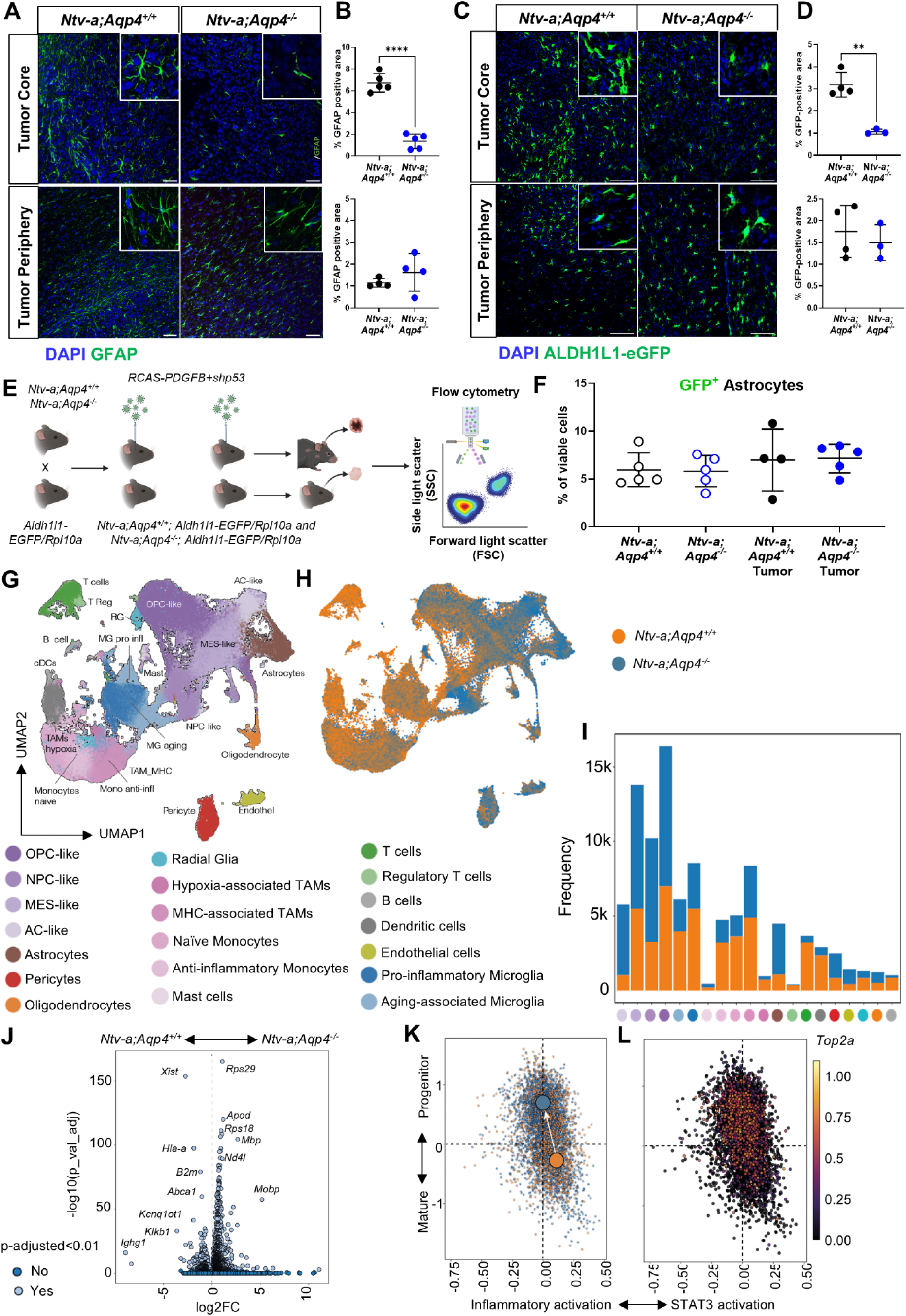
Loss of *Aqp4* results in decreased astrocyte coverage within the tumor core and an astrocytic shift to a less reactive, progenitor-like state. **A**, Representative images showing GFAP staining (green) within the tumor core and periphery of *Ntv-a;Aqp4^+/+^* and *Ntv-a;Aqp4^-/-^* mice bearing PDGFB mGBM. DAPI was used as a nuclear stain. Images were taken on the Leica SP8 confocal microscope. Scale bar=20 μm. B, Quantification of GFAP-positive area indicates a decrease in coverage within the tumor core of mice lacking AQP4 but not the tumor periphery. Two-tailed Student’s t-test, ****p<0.0001. C, Representative images showing GFP staining (green) within the tumor core and periphery of *Ntv-a;Aqp4^+/+^;Aldh1l1-eGFP/Rpl10a* and *Ntv-a;Aqp4^-/-^;Aldh1l1-eGFP/Rpl10a* mice bearing PDGFB mGBM. DAPI was used as a nuclear stain. Images were taken on the Leica SP8 confocal microscope. Scale bar=100 μm. D, Quantification of GFP-positive area indicates a decrease in coverage within the tumor core of mice lacking AQP4 but not the tumor periphery. E, Schematic illustration of the generation of *Ntv-a;Aqp4^+/+^;Aldh1l1-eGFP/Rpl10a* and *Ntv-a;Aqp4^-/-^;Aldh1l1-eGFP/Rpl10a* mice, followed by tumor generation and flow cytometry to quantify the GFP^+^ astrocyte population. F, Graph comparing the population of GFP^+^ astrocytes in naïve and tumor-bearing *Ntv-a;Aqp4^+/+^* and *Ntv-a;Aqp4^-/-^* mice as a percentage of the total viable cells. G, UMAP dimensionality reduction and cluster annotation of scRNA-seq data acquired from n=4 *Ntv-a;Aqp4^+/+^*and n=3 *Ntv-a;Aqp4^-/-^* PDGFB mGBM tumors. H, UMAP representation of differential abundance of *Ntv-a;Aqp4^+/+^* (orange) and *Ntv-a;Aqp4^-/-^* (blue) samples within the scRNA-seq data. I, Graph showing frequency of various clusters identifed by dimensionality reduction between *Ntv-a;Aqp4^+/+^* (orange) and *Ntv-a;Aqp4^-/-^* (blue) samples. J, Volcano plot showing the differentially expressed genes within the astrocyte subcluster in *Ntv-a;Aqp4^+/+^* and *Ntv-a;Aqp4^-/-^* samples. K, Four-dimensional state plot of the astrocyte subcluster in *Ntv-a;Aqp4^+/+^*(orange) and *Ntv-a;Aqp4^-/-^* (blue) samples, where the X-axis differentiates between inflammatory and STAT3-mediated activation of astrocytes and the Y-axis differentiates between mature and progenitor astrocyte states. **L**, Four-dimensional state plot of the astrocyte subcluster from **K** with *Top2a* expression overlay.

**Fig. 5:**
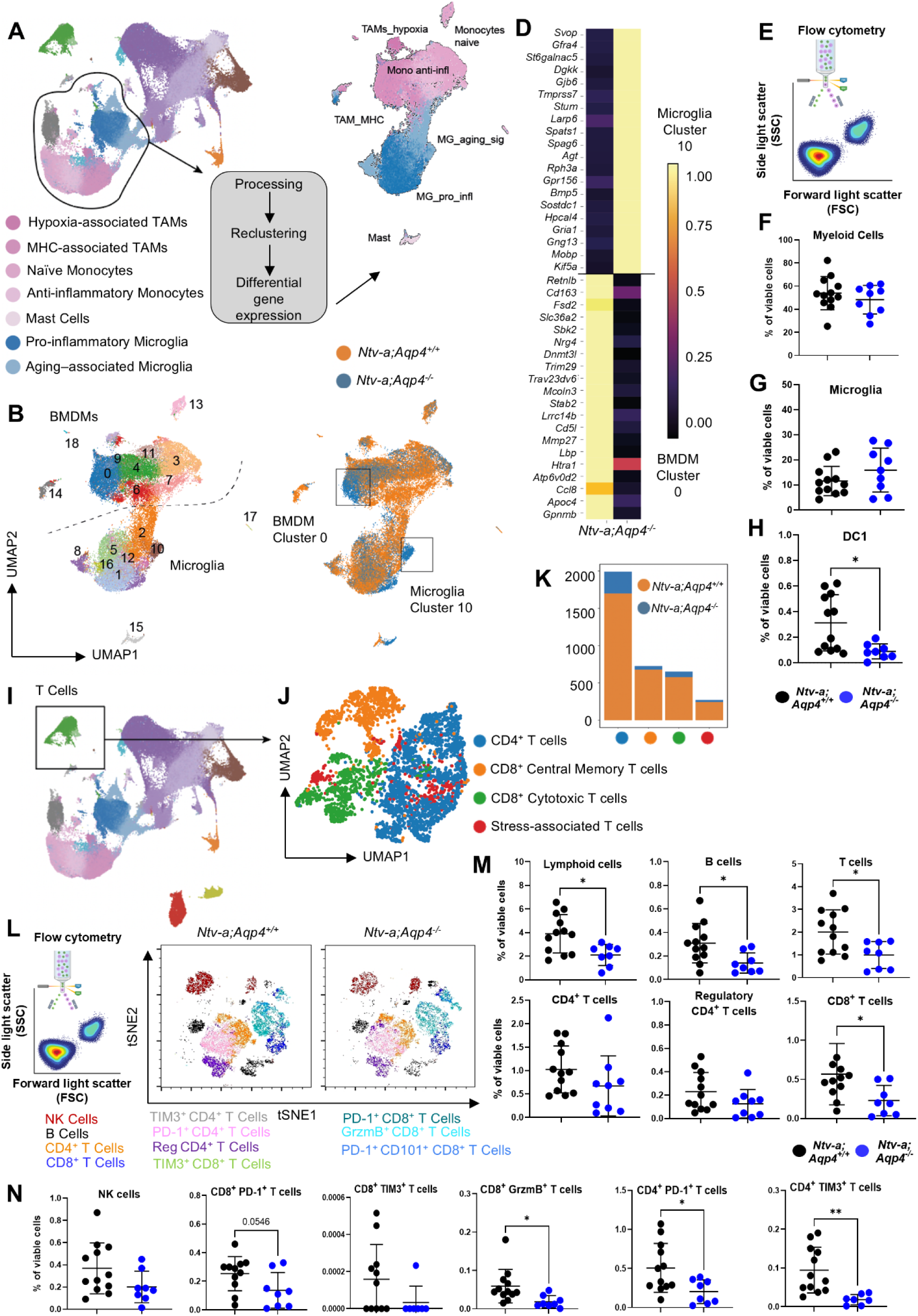
Loss of *Aqp4* results in changes within the myeloid microenvironment as well as a decrease in T cells in PDGFB mGBM. **A**, UMAP dimensionality reduction of scRNA-seq data from the myeloid compartment and representation of the cell types. **B**, Sub-clustering and annotation of bone marrow derived myeloid cells and microglia. **C**, UMAP representation of *Ntv-a;Aqp4^+/+^* (orange) and *Ntv-a;Aqp4^-/-^* (blue) samples within the myeloid compartment of scRNA-seq data. **D**, List of differentially enriched genes in Cluster 0 from the BMDM compartment and Cluster 10 from the Microglia compartment within *Ntv-a;Aqp4^-/-^* samples compared to *Ntv-a;Aqp4^+/+^* samples. **E, F, G and H**, Schematic illustration of flow cytometry performed on tumors from *Ntv-a;Aqp4^+/+^* and *Ntv-a;Aqp4^-/-^* mice and graphs comparing the population of myeloid cells, microglia and dendritic cells type I between the two genotypes as a percentage of total viable cells. Two-tailed Student’s t-test, *p<0.05. **I and J**, UMAP dimensionality reduction of scRNA-seq data from the T cell compartment and representation of the cell types. **K**, Frequency of T cell clusters as identified in **J** between *Ntv-a;Aqp4^+/+^* and *Ntv-a;Aqp4^-/-^* tumor-bearing mice. **L**, tSNE plot showing proportions of T cell subpopulations in *Ntv-a;Aqp4^+/+^*and *Ntv-a;Aqp4^-/-^* mice bearing PDGFB mGBM. **M and N**, Graphs comparing the lymphoid populations in *Ntv-a;Aqp4^+/+^* and *Ntv-a;Aqp4^-/-^* mice as a percentage of total viable cells. Two-tailed Student’s t-test, *p<0.05; **p<0.01.

### Loss of *Aqp4* leads to a phenotypic shift in the myeloid compartment and a decrease in lymphoid cells in tumors

In addition to the astrocyte subcluster, our scRNA-seq analyses also revealed decreases in the myeloid and lymphoid subclusters in *Aqp4-*deficient PDGFB mGBM (Fig. 4I). Moreover, reactive astrocytes are known to recruit and interact with several other cell types in pathological conditions including GBM (30, 68, 69), such as microglia and BMDMs (collectively called tumor-associated macrophages or TAMs), which make up approximately 40% of the GBM tumor mass (70), as well as T cells (71). We next probed changes in the immune microenvironment of *Aqp4-*proficient and *Aqp4-*deficient mice. We performed spectral flow cytometry on PDGFB mGBM tumors using a previously-established panel of immune markers and a gating strategy that allowed for accurate separation of various myeloid and lymphoid populations in GBM (55, 72, 73). There were no differences between *Aqp4*-proficient and *Aqp4*-deficient tumors in the relative proportion of myeloid subpopulations (Fig. 5E-G and Supp. Fig. 7A-O), except for a decrease in dendritic cells type I (DC1) in *Aqp4*-deficient mice (Fig. 5H). To further dissect phenotypic changes in the myeloid compartment, we performed re-clustering of myeloid cells from our scRNA-seq dataset, which allowed for a more refined definition of distinct subpopulations and their distribution across *Aqp4*-proficient and *Aqp4*-deficient tumors (Fig. 5A&B). This analysis revealed two subgroups (BMDM cluster 0 and Microglia cluster 10) that were enriched in *Aqp4*-deficient tumors (Fig. 5C). Differential gene expression analysis identified the cluster 0 BMDM subpopulation characterized by expression of genes associated with tissue remodeling, immune modulation, metabolic adaptation, matrix degradation, and lysosomal activity. Interestingly, many of the differentially-expressed genes such as *Cd163, Cd5l*, *Gpnmb,* and *Ccl8* were associated with an immunosuppressive and pro-tumorigenic macrophage phenotype (74–77) (Fig. 5D). We also performed linear regression analysis of CD15-CD31 positive areas and AQP4-CD163 positive areas (Extended Data Fig. 1A,G&H) from human GBM sections stained using the Orion^™^ multiplex IF platform. Higher CD15⁺ myeloid infiltration correlated with increased vascular density, whereas CD163⁺ TAM enrichment showed an inverse correlation with AQP4 expression. Notably, anti-VEGF treatment has previously been shown to reduce myeloid infiltration in murine GBM, consistent with our finding that myeloid influx positively correlates with vascular density. (78). In parallel, we identified a distinct microglial subpopulation (cluster 10) characterized by genes such as *Sostdc1*, *Bmp5*, and *Gria1*, indicative of a potential role in neuromodulation and BMP-signaling pathways (79, 80). This subpopulation was additionally linked to neurodevelopmental processes, extracellular signaling, and tissue remodeling, suggesting a contribution to microglia-mediated neuroprotection and homeostasis within the tumor microenvironment. The differential enrichment of these myeloid subpopulations in *Aqp4*-deficient mice indicates that *Aqp4* loss correlates with changes in immune cell composition, shifting TAM function towards a state associated with immunosuppression and homeostasis. To investigate the lymphoid population, we isolated T cells from our scRNA-seq data and performed re-clustering followed by automated cell type annotation to profile T-cell subgroups within *Aqp4*-deficient mice bearing PDGFB mGBM (Fig. 5I&J). We found that T cells demonstrated an overall lower abundance in *Aqp4*-deficient mice as compared to *Aqp4*-proficient mice. The most significant decrease was observed in the cytotoxic CD8^+^ and stress-associated T-cell subpopulations (Fig. 5K). Because of the low numbers of T cells in *Aqp4*-deficient mice, differential gene expression did not identify any significant changes. We also employed lymphoid cell-specific markers to dissect the various subpopulations in our spectral flow cytometry data (Fig. 5L and Supp. Fig. 7P&Q). Consistent with our scRNA-seq findings, there was a decrease in the proportion of lymphoid cells in *Aqp4*-deficient mice compared to their wild-type counterparts. This decrease was significant in B cells and T cells (Fig. 5M). To assess the activation and functional states of CD4^+^ and CD8⁺ T cells, we quantified the proportion of cells expressing the inhibitory receptors PD-1 (Programmed cell death protein-1 or CD279) and TIM-3 (T-cell immunoglobulin and mucin domain-3) as well as the cytotoxic effector molecule GrzmB (Granzyme B). Compared to *Aqp4*-proficient mice, *Aqp4*-deficient mice displayed a trend towards decreased proportions of PD-1⁺ and TIM-3⁺ CD8⁺ T cells (Fig. 5N), although the expression of these markers did not change (Supp. Fig. 7T&U), indicating lower levels of tumor antigen-driven activation and exhaustion. The frequency of GrzmB⁺ CD8⁺ T cells was also significantly reduced, suggesting impaired cytotoxic potential (Fig. 5N). Within the CD4⁺ T cell population, the frequencies of PD-1⁺ and TIM-3⁺ CD4⁺ subsets were significantly lower in *Aqp4*-deficient mice, again without changes in expression of these markers (Fig. 5M&N and Supp. Fig. 7R&S). These findings indicate a reduction in both CD8⁺ and CD4⁺ T-cell infiltration and activation in *Aqp4*-deficient mice, and, together with our findings in the myeloid subset, suggest a weakened immune response.

### Loss of *Aqp4* reshapes the tumor–brain interface and dampens tumor-associated inflammation

To investigate the role of AQP4 in shaping the spatial architecture of PDGFB mGBM, we generated whole-section Visium maps from *Aqp4*-proficient and deficient tumors (n=4 per genotype). Each section yielded a median of 4.9 × 10³ genes and 1.8 × 10⁴ unique molecules per spot, providing a total of 20,412 spatially-indexed transcriptomes across the cohort (Fig. 6A). We first integrated our scRNA-seq dataset (Fig. 4) with Visium data using cell2location (81) and CytoSpace (82), assigning 27 cell states—including malignant, immune, and glial populations—to each spot (Fig. 6B&C). These annotations were then used as input for NePSTA (83), a graph deep-learning framework that tiles non-overlapping “cellular ecosystems” while simultaneously predicting both the genotype and the region of origin (tumor vs. normal brain) for each ecosystem (Fig. 6D). Stratified five-fold cross-validation achieved an accuracy of 0.98 (95% CI 0.96–1.00) and an F1-score of 0.99, demonstrating that *Aqp4* loss imprints a distinct spatial signature on the tumor microenvironment. Plotting the logit score for each of the 1,669 ecosystems against genotype and regional probability revealed two well-separated clusters (Fig. 6E). Notably, *Aqp4*-deficient ecosystems showed enrichment of neuronal populations and astrocytes within the tumor core as well as alterations in the peritumoral area, leading to a shift in the regional prediction capacity towards non-tumoral regions (p = 0.0321). This suggests that *Aqp4* loss affects not only the tumor, but also its surrounding microenvironment. Ecosystems enriched in *Aqp4*-proficient tumors were characterized by abundant T cells, dendritic cells, and inflammatory myeloid populations (TAM MHC, MG pro-inflammatory), whereas *Aqp4*-deficient ecosystems were dominated by pericytes, astrocytes, oligodendrocyte-lineage cells, and NPC-/OPC-like malignant cells (Fig. 6F), consistent with our scRNA-seq dataset. To further dissect the molecular programs driving these patterns, we aggregated the top 50 ecosystems (highest F1 score per genotype) into pseudo-bulk profiles and performed differential expression followed by gene set enrichment analysis. *Aqp4-*deficient niches preferentially expressed neuronal and synaptic pathways—including “synaptic vesicle priming,” “glutamatergic synapse,” and “regulation of synaptic plasticity”—whereas *Aqp4*-proficient niches upregulated cytokine- and antigen-processing modules (Fig. 6G). These findings support the notion that elevated intracranial pressure and edema can trigger axonal injury, thereby activating neuronal and synaptic pathways that have been shown to promote tumor growth (57, 84, 85). Overall, these results suggest that *Aqp4* loss shifts the tumor microenvironment away from an inflamed, immune-rich state toward one resembling normal brain tissue, in line with the NePSTA prediction scores. It is plausible that mechanical constraints imposed by the skull, coupled with elevated intracranial pressure, induce axonal injury, which may contribute to reduced survival, as recently demonstrated Cell–cell neighborhood analysis centered on astrocytes further highlighted genotype-specific rewiring (Fig. 6H&I). In *Aqp4*-proficient tumors, astrocytes neighbored AC-like tumor cells, radial glia-like (RG) cells, and activated T cells. In contrast, *Aqp4*-deficient tumors displayed astrocytic neighborhoods dominated by oligodendrocyte precursor cells (OPCs), mature oligodendrocytes, microglia, and NPC-/OPC(85)-like tumor cells. Quantitative analysis confirmed a two-fold reduction in astrocyte–T cell adjacency (adj. p < 0.01, two-sided Wilcoxon) and a three-fold increase in astrocyte–OPC contacts (adj. p < 0.001). Together, these data demonstrate that *Aqp4* loss attenuates anti-tumor immunity and promotes a more ‘brain-like’ synaptogenesis-oriented ecosystem at the tumor margin. Given the observed enrichment of astrocytes at both the tumor core and margin, we next assessed astrocytic gene expression. Cell-type-specific gene expression estimation (86) was performed using the astrocytic enrichment score, followed by enrichment analysis of TAA subtypes (inflammatory-activated vs. STAT3-activated). In *Aqp4*-proficient tumors, astrocytes at the tumor margin predominantly displayed reactive phenotypes, consistent with previous reports (Fig.6J). In contrast, *Aqp4*-deficient tumors showed enrichment of inflammatory astrocytes within the tumor core and STAT3-activated astrocytes at the tumor margin —findings that align closely with our scRNA-seq analysis.

**Fig. 6:**
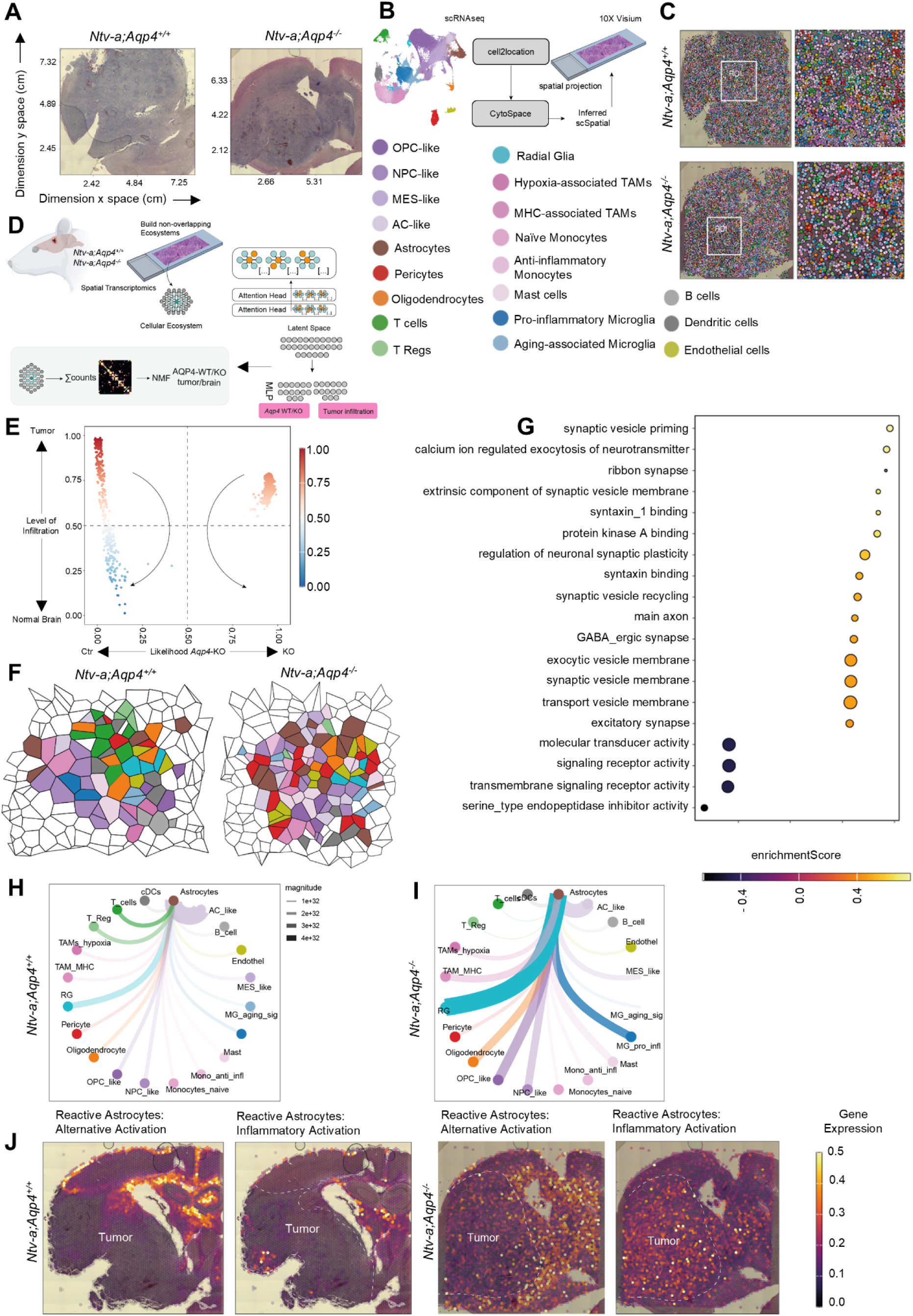
Loss of *Aqp4* affects spatial distribution of tumor-associated astrocytes and significantly impacts their interactions with other cell types in the tumor microenvironment. **A**, Bright-field overview of Visium sections from *Ntv-a;Aqp4^+/+^* (left) and *Ntv-a;Aqp4^-/-^* (right) PDGFB mGBM brains showing tumor (dark) and surrounding tissue. B&C, Integration workflow: single-cell RNA-seq reference mapped onto Visium spots with *cell2location* and CytoSpace; representative section colored by 27 cell states (legend). **D**, Schematic of the NePSTA pipeline. Spots were tiled into non-overlapping cellular ecosystems (hop=6); a graph neural network classified each ecosystem by genotype and tumor vs non-tumor regions. **E**, Ecosystem ranking plot. Each point is an ecosystem (n=1,669); x-axis shows logit score for *Ntv-a;Aqp4^-/-^*, y-axis shows the logit score of tumor vs non-tumor origin. **F**, Representative cellular ecosystem tilings for *Ntv-a;Aqp4^+/+^* and *Ntv-a;Aqp4^-/-^* sections; each polygon indicates one ecosystem, colored by dominant cell type. **G**, Differential pseudo-bulk gene-set enrichment comparing the top-50 ecosystems between *Ntv-a;Aqp4^-/-^* and *Ntv-a;Aqp4^+/+^* samples. Dot size represents the -log10 adj. p-value ; colors indicate NES (normalized enrichment score). **H&I**, Neighborhood analysis centered on astrocytes with Circos plots depicting the frequency-weighted edges between astrocytes (center) and neighboring cell states in *Ntv-a;Aqp4^+/+^* (left) and *Ntv-a;Aqp4^-/-^* (right) tumors. Edge width reflects interaction magnitude (see legend). **J**, Changes in reactive astrocyte localization with astrocytic rim around the tumors in *Ntv-a;Aqp4^+/+^* samples and enhanced reactive astrocytes within the tumor core of *Ntv-a;Aqp4^-/-^* samples.

### CD4^+^ and CD8^+^ T-cell depletion in *Aqp4*-proficient and *Aqp4*-deficient tumor-bearing mice has no effect on survival

Given our finding that *Aqp4*-deficient tumors had reduced lymphoid cell infiltration (Fig. 5I-N), we next considered whether AQP4 directly influences T-cell biology. AQP4 has been reported to be expressed on both CD4^+^ and CD8^+^ T cells (87), and functional studies demonstrate that it is required for TCR-mediated activation via actin cytoskeleton remodeling (88), as well as for chemokine production and lymphocyte trafficking (89). These observations, together with the diminished T-cell presence in *Aqp4*-deficient tumor-bearing mice, prompted us to investigate whether *Aqp4* loss influences the contribution of T cells to the reduced survival of *Aqp4*-deficient compared to *Aqp-4* proficient tumor-bearing mice. We generated PDGFB mGBM in *Aqp4*-proficient and *Aqp4*-deficient mice and performed CD4^+^ and CD8^+^ T-cell depletion in two separate cohorts. In the CD8 depletion cohort, 200 μg of anti-CD8 antibody or IgG2b control was administered every 4 days starting at Day 20 post-injection and continuing until endpoint signs developed (Fig. 7A). In the CD4 depletion cohort, 300 µg of anti-CD4 antibody or IgG2b control was initially administered at Day 22 post-injection and again two days later, followed by treatment every 7 days until endpoint signs developed (Fig. 7D). We also harvested tumors and blood at endpoint for flow cytometry analysis. In the CD8 depletion cohort, we did not observe any differences in survival between anti-CD8 and IgG2b-treated *Aqp4*-proficient (m.s 44 vs. 44 days) and *Aqp4*-deficient (m.s 42 vs. 43 days) mice (Fig. 7B&C). Similarly, in the CD4 depletion cohort, anti-CD4 treatment did not have a significant impact on survival in *Aqp4*-proficient (m.s 48 vs. 55 days) and *Aqp4*-deficient mice (m.s 44 vs. 51 days) (Fig. 7E&F). While we confirmed depletion of the CD8^+^ and CD4^+^ T-cell populations in both experiments (Supp. Figs. 8C&D,9A&B,10A&B), we did not observe any differences in other immune cell populations in the tumor or blood by flow cytometry (Supp. Figs. 8-10). Together, these findings suggest that AQP4 status does not shape T cell–mediated control of tumor growth in this model, and that adaptive immunity has a limited role in determining disease outcome. The lack of efficacy of CD8^+^ or CD4^+^ T-cell depletion in *Aqp4*-proficient mice can be attributed to their very low abundance in GBM.

**Fig.7.**
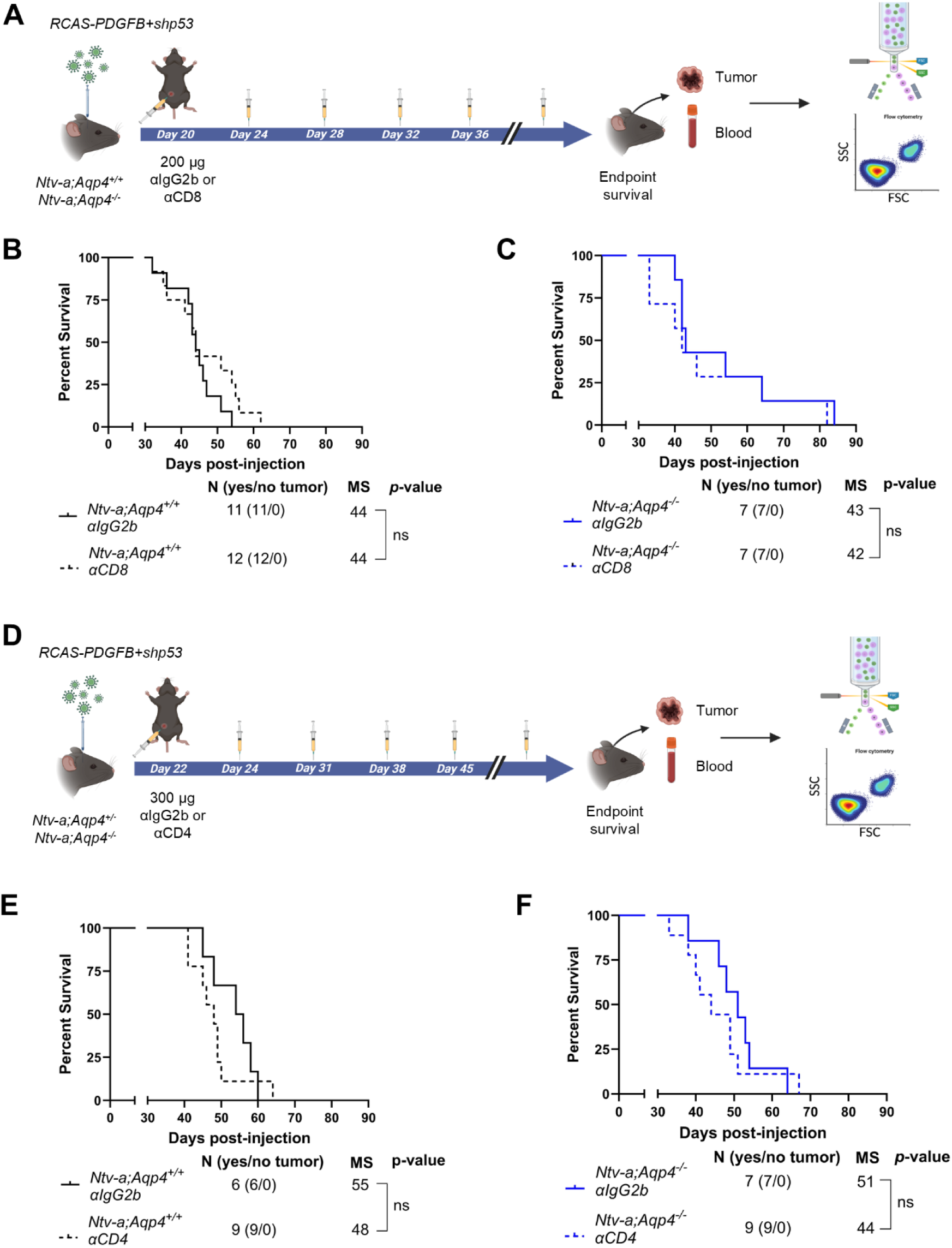
Depletion of CD8^+^ and CD4^+^ T cells in *Ntv-a;Aqp4^+/+^* and *Ntv-a;Aqp4^-/-^* mice bearing PDGFB-driven tumors has no effect on survival. **A**, Schematic showing CD8+ T cell depletion in *Ntv-a;Aqp4^+/+^* and *Ntv-a;Aqp4^-/-^* mice bearing PDGFB mGBM, followed by blood/tumor harvest and flow cytometry. **B & C**, Kaplan-Meier survival curves comparing median survival between anti-CD8 treated and IgG2b-treated mice in *Ntv-a;Aqp4^+/+^* (left) and *Ntv-a;Aqp4^-/-^* (right) groups. Curves were compared by log-rank (Mantel-Cox) test. **D**, Schematic showing CD4+ T cell depletion in *Ntv-a;Aqp4^+/+^* and *Ntv-a;Aqp4^-/-^* mice bearing PDGFB mGBM, followed by blood/tumor harvest and flow cytometry. **E & F,** Kaplan-Meier survival curves comparing median survival between anti-CD4 treated and IgG2b-treated mice in *Ntv-a;Aqp4^+/+^* (left) and *Ntv-a;Aqp4^-/-^* (right) groups. Curves were compared by log-rank (Mantel-Cox) test.

### Anti-VEGF treatment reduces vessel leakage and increases *Aqp4* expression in tumor-bearing mice

Since adaptive immune responses were unlikely to contribute significantly to observed differences between *Aqp4*-proficient and *Aqp4*-deficient tumors (Fig. 7), we turned our attention to the vasculature within the tumor microenvironment. Angiogenesis and microvascular proliferation are well-established hallmarks of GBM (90). AQP4 has been implicated in maintaining BBB integrity during development and in a murine model of intracerebral hemorrhage (91–93), but has not been investigated in this regard in GBM. Thus, we first investigated the effect of *Aqp4* loss on Hoescht dye leakage as a marker of vessel leakiness. *Aqp4*-deficient tumor-bearing mice had higher levels of vessel leakage compared to *Aqp4*-proficient mice (Fig. 8G). Anti-angiogenic therapy with bevacizumab (Avastin) is FDA-approved for recurrent GBM, but while it transiently improves progression-free survival and edema, it does not provide an overall survival benefit (10–12). However, given bevacizumab’s well-recognized anti-angiogenic effect, we sought to investigate whether anti-angiogenic treatment could alleviate the increased vessel leakage observed in the *Aqp4*-deficient tumor-bearing mice. We treated PDGFB mGBM-bearing *Aqp4*-proficient and *Aqp4*-deficient mice with a neutralizing anti-VEGFA antibody, B20-4.1.1 (which recognizes both human and murine VEGFA), followed by assessment of survival and vessel leakage (Fig. 8A). We did not observe a significant survival difference between B20-4.1.1-treated and control-treated mice in either the *Aqp4*-proficient or *Aqp4*-deficient cohorts (Fig. 8B&C). Although there were trends toward reduced edema in the B20-4.1.1 cohorts for both genotypes, statistical significance was not reached (Supp. Fig. 11A&B). These results differ from our previously-published work using GBM generated in *Cdkn2a* germline-deficient mice, where we observed a significant reduction in edema content and improved survival of tumor-bearing mice, as measured using more sensitive techniques, such as T2-weighted MRI (13). Another contributing factor to this difference can be the fact that, in *Cdkn2a* germline knockout mice, *Cdkn2a* loss occurs not only in tumor cells, but also in endothelial cells, which may contribute to increased proliferation and angiogenesis in these tumors. Therefore, the effects of anti-VEGFA blockade can be more pronounced in this model. We then compared vessel leakiness in B20-4.1.1-treated and control-treated mice within the two genotypes. As we have previously reported, B20-4.1.1-treated *Aqp4*-proficient mice had decreased vessel leakage compared to control-treated mice (13), while this decrease was not observed in the *Aqp4*-deficient group (Fig. 8D-F). To assess the effect of anti-VEGF blockade on brain aquaporin levels, we ran a qPCR for *Aqp4* and *Aqp1*. Our results showed that B20-4.1.1-treated mice specifically upregulated *Aqp4* expression, while *Aqp1*, which is associated with endothelial cells in the brain vasculature (94), was not upregulated (Fig. 8H&I). This, along with the leakage assay results, indicates that the effect of anti-VEGF blockade on vascular leakiness is AQP4-dependent. We further studied the characteristics of the tumor-associated vasculature in *Aqp4*-deficient mice through blood vessel (CD31) and pericyte (alpha-smooth muscle actin; alpha-SMA) staining. The average size of CD31-positive blood vessels and the CD31-positive area did not differ between *Aqp4*-proficient and *Aqp4*-deficient mice (Fig. 8J&K and Supp. Fig. 11C). Similarly, alpha-SMA did not differ in terms of positive area and staining intensity between the two groups (Fig. 8L&M and Supp. Fig. 11D). Thus, while *Aqp4* loss was found to impact astrocyte coverage within the tumor (Fig. 4), it did not have significant effects on other cell types forming the BBB such as endothelial cells and pericytes. However, our results indicate a previously-unknown effect of *Aqp4* loss on promoting vascular leakiness in GBM, and that the protective effect of anti-VEGF treatment depends on AQP4, as reduced permeability was observed only in *Aqp4*-proficient but not in *Aqp4*-deficient mice.

**Fig. 8.**
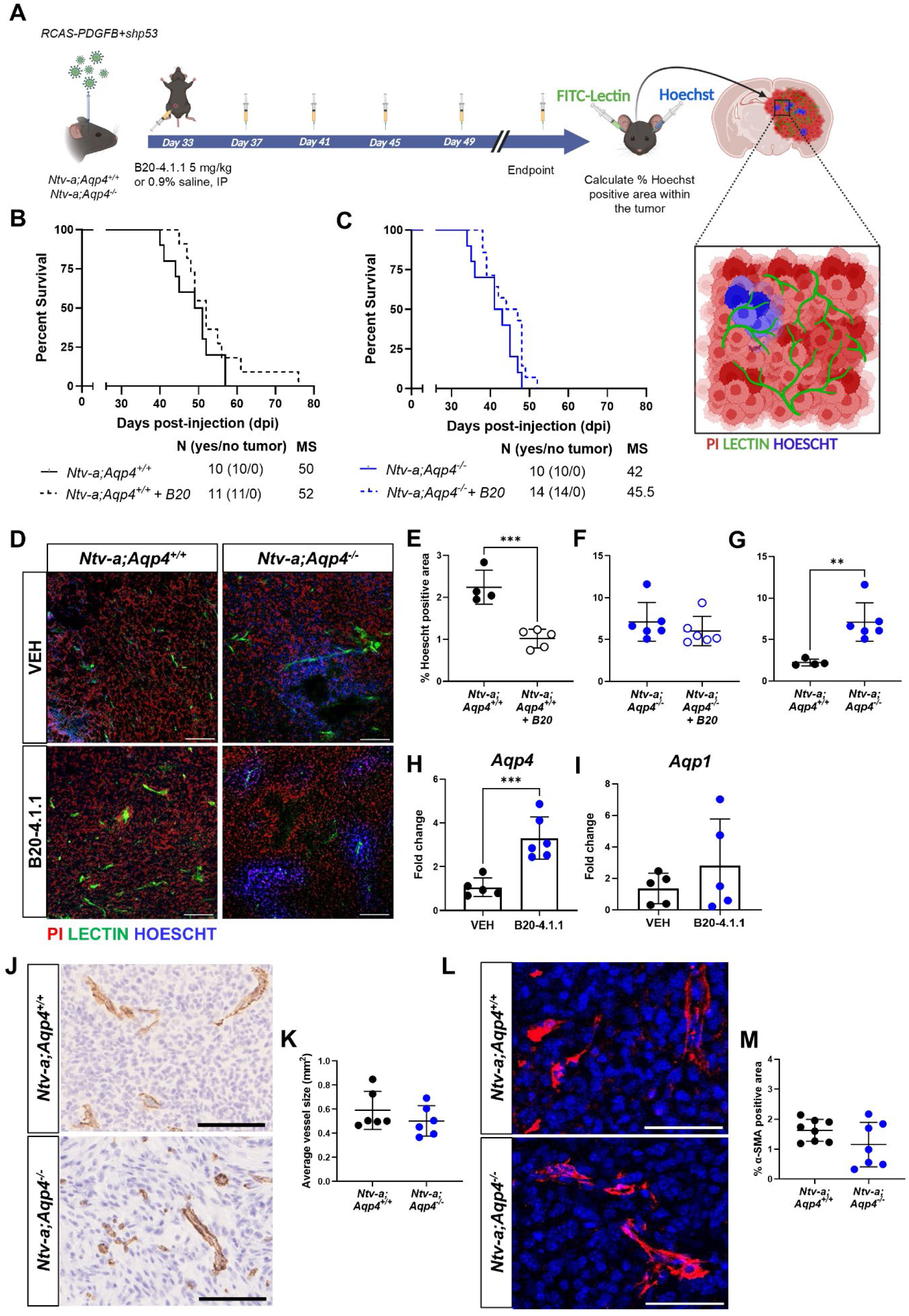
Anti-VEGF treatment reduces vessel leakage and increases *Aqp4* expression in wild-type tumor-bearing mice. **A**, Schematic illustration of treatment with B20-4.1.1, a murine analog of Bevacizumab, in PDGFB mGBM-bearing mice followed by vessel leakage assay. **B and C**, Kaplan-Meier survival curves demonstrating no significant difference in survival between B20-4.1.1 and control-treated *Ntv-a;Aqp4^+/+^* and *Ntv-a;Aqp4^-/-^* mice bearing PDGFB mGBM. Curves were compared using the log-rank (Mantel Cox) test. **D**, Representative images showing vessel leakage marked by Hoescht staining (blue), blood vessels marked by FITC-Lectin (green) and tumor cells marked by propidium iodide (red). All images were taken on a Leica SP8 confocal microscope. Scale bar=100 μm. **E and F**, Graphs showing percentage of Hoescht positive area compared between B20-4.1.1 and control-treated *Ntv-a;Aqp4^+/+^*and *Ntv-a;Aqp4^-/-^* mice bearing PDGFB mGBM. Two-tailed Student’s t-test, ***p<0.001. **G**, Graph showing percentage of Hoescht positive area compared between control-treated *Ntv-a;Aqp4^+/+^* and *Ntv-a;Aqp4^-/-^*mice bearing PDGFB mGBM. Two-tailed Student’s t-test, **p<0.01. **H and I**, Graphs showing fold change in *Aqp4* and *Aqp1* gene expression respectively over houskeeping gene *β-actin* compared between B20-4.1.1 and control-treated *Ntv-a;Aqp4^+/+^* tumor-bearing mice. Two-tailed Student’s t-test, ***p<0.001**. J and K**, Representative images (left) showing IHC staining for CD31 and quantification (right) of average blood vessel size (in mm2) in CD31 positive areas from *Ntv-a;Aqp4^+/+^* (n=6) and *Ntv-a;Aqp4^-/-^* (n=6) mice bearing PDGFB mGBM. Two-tailed Student’s t-test. Scale bar=100 μm**. L and M**, Representative images (left) showing IF staining for alpha-SMA and quantification (right) of alpha-SMA positive area (percentage) from *Ntv-a;Aqp4^+/+^* (n=7) and *Ntv-a;Aqp4^-/-^*(n=7) mice bearing PDGFB mGBM. Two-tailed Student’s t-test. Scale bar=50 μm

## Discussion

Cerebral edema remains one of the most clinically important problems, yet it remains understudied in GBM. Here, using multi-regional human GBM biopsies, murine models of *de novo* GBM, and integrative single-cell and spatial profiling, we identify the water channel AQP4 as a central regulator of edema clearance and uncover broader roles in astrocyte maturation, vascular integrity, and immune composition and phenotype within the tumor microenvironment.

AQP4 is concentrated at astrocytic endfeet along perivascular and subpial surfaces and is the main water channel in the central nervous system (CNS). It enables rapid water exchange between brain parenchyma, cerebrospinal fluid, and blood (22). Early studies have shown that AQP4 exhibits context-dependent, opposing functions in cerebral edema. Under conditions where the BBB is intact, AQP4 activity contributes to the development of cytotoxic edema. By contrast, when BBB integrity is compromised, AQP4 facilitates the clearance of vasogenic edema (23, 24). Despite extensive work in other models, including stroke, injury, and one study using a metastatic brain tumor model of B16F10 melanoma, there are currently no published studies that directly assess the distinct roles of AQP4 in murine GBM models. The RCAS/*tv-a* model (49, 50) enabled us to test whether the inverse association between AQP4 expression levels and edema observed in patients extends beyond correlation. *Aqp4-*deficient GBM-bearing mice exhibited increased brain water content, shortened survival, and smaller tumor volumes, indicating that excess water accumulation, rather than tumor size, drives the survival deficit.

Although AQP4 is linked to GBM-associated edema, its cellular origin remains controversial. Early histological studies reported that AQP4 expression increases with glioma grade and is often distributed ubiquitously across the tumor (25–27). Additionally, using human and rat glioma cell line-based models it was suggested that tumor cells express AQP4 (28, 29). Our *in situ* hybridization and multiplex immunofluorescence data resolve this long-standing uncertainty by showing that AQP4 localizes mainly to reactive astrocytes, rather than tumor cells, in both human and murine GBM. Although AQP4 levels are increased in human GBM, its redistribution was suggested to impair its function (95), which we confirmed in human and provide direct evidence of functional impairment in our murine models. In *Aqp4*-deficient tumors, astrocyte coverage within the tumor core was markedly reduced, although the overall number of astrocytes remained unchanged. scRNAseq data revealed that astrocytes lacking *Aqp4* adopted an immature, progenitor-like transcriptional program, characterized by upregulation of *Top2a* and downregulation of antigen-presentation and immune-response genes. This immature transcriptional state closely resembles fetal or de-differentiated astrocytes previously described at the tumor margin in human GBM (30, 67), suggesting that AQP4 is a molecular determinant of the tumor-associated reactive phenotype. Spatial transcriptomics revealed that astrocytes in WT tumors were spatially associated with T cells, radial glia-like cells, and AC-like tumor cells, whereas those in *Aqp4*-deficient tumors preferentially neighbored oligodendrocyte precursor cells and neuronal-like populations. The decrease in astrocyte–T cell adjacency and the emergence of oligodendrocyte and synaptic signatures within the tumor core indicate a disruption of the normal glial-immune interface. Although glial scar-like astrocytic barriers have classically been described in brain injury or pathology (96), our data suggest that when AQP4 is lost, astrocytes fail to maintain these boundaries, potentially allowing for abnormal synaptic integration that could promote tumor invasion. Tumors lacking *Aqp4* were depleted of lymphoid cells and enriched for immunosuppressive, tissue-remodeling TAMs. In contrast, *Aqp4*-proficient tumors contained higher proportions of MHC-enriched TAMs, dendritic cells, and activated T cells, suggesting a microenvironment more conducive to immune surveillance. Although depletion of lowly abundant CD4⁺ or CD8⁺ T cells did not affect survival - consistent with the limited role of adaptive immunity in GBM (97) - the reduced abundance and activation of lymphoid cells in *Aqp4*-deficient tumors indicates that AQP4 sustains some degree of immune competence in GBM. The absence of immune-rich niches in *Aqp4*-deficient tumors and the dominance of neuronal and synaptic gene programs indicates a global shift toward an immune-quiescent, “brain-like” state. These immunological changes may exacerbate edema and tissue damage by impairing phagocytic clearance, but the mechanisms remain to be determined. *Aqp4*-deficient tumors displayed pronounced vascular leakiness, suggesting that AQP4 also plays a role in vascular integrity. Notably, treatment with an anti-VEGFA antibody reduced vessel leakage only in *Aqp4*-proficient mice and upregulated *Aqp4* expression, suggesting that AQP4 is required for the full effect of VEGF blockade. This relationship also suggests that VEGF inhibition may restore vascular integrity by enhancing astrocyte-endothelial cell interactions through AQP4.

Our findings extend this view by showing that this upregulation is closely associated with astrocyte polarization and organization of immune-glial interfaces, suggesting that its role in GBM extends beyond edema resolution to the maintenance of tissue architecture. While pharmacological inhibition of AQP4 surface localization has been shown to mitigate acute cytotoxic swelling (98), our data suggest that in the chronically disrupted GBM environment, strategies aimed at restoring or enhancing AQP4 at astrocytic endfeet may instead be beneficial.

## Methods

### Mice

All experimental procedures were approved by the IACUC (Institutional Animal Care and Use Committee) at the Icahn School of Medicine at Mount Sinai (Protocol #2019-00619 and #2014-0229) and Perelman School of Medicine, University of Pennsylvania (Protocol #807737). Age- and sex-matched mice between 6 and 12 weeks old were used for all experiments. All mice were in a C57BL/6J background and housed in a climate-controlled, pathogen-free facility with access to food and water *ad libitum* under a 12-hour light/dark cycle. Genotyping of all mice used for experiments was performed using Transnetyx (Cordova TN). The aquaporin-4 (*Aqp4*) germline knockout mice were obtained from Dr. Alan Verkman (99) (University of California, San Francisco) and crossed to *Ntv-a* transgenic mice to obtain *Ntv-a; Aqp4*^-/-^ mice.

### Generation of *de novo* murine GBM

Briefly, chicken fibroblast cells (DF-1; ATCC #CRL-3586) were grown at 39°C, 5% CO_2_ according to supplier instructions. Cells were transfected with RCAS-*hPDGFB*-*HA*, RCAS-*hPDGFA*-myc, RCAS-shRNA-*p53*-RFP, RCAS-shRNA-*Nf1*, and RCAS-shRNA-*Pten*-RFP using the Fugene 6 transfection kit (Promega #E5911). To generate tumors, mice were anesthetized using ketamine (100 mg/kg) and xylazine (10 mg/kg) and approximately 40,000 transfected DF-1 cells were stereotactically delivered using a Hamilton syringe with a 30-gauge needle in 1 µl neurobasal medium per mouse. A mixture of equal parts RCAS-*hPDGFB*-*HA* and RCAS-shRNA-*p53*-RFP was injected to generate *PDGFB*-driven GBM in the right frontal striatum (coordinates: AP −1.5 mm and right −0.5 mm from bregma; depth −1.5 mm from the dura surface). Similarly, a mixture of equal parts RCAS-*hPDGFA*-myc, RCAS-shRNA-*p53*-RFP, RCAS-shRNA-*Nf1*, and RCAS-shRNA-*Pten*-RFP was injected to generate *NF1*-silenced GBM (coordinates: AP −0.0 mm and right −0.5 mm from bregma; depth −1.5 mm from the dura surface). For EGFRvIII mGBM, a combination of RCAS-*Cre*, RCAS-*TAZ,* and RCAS-*shp53* was injected in *Cdkn2a^−/−^*;*EGFRvIII^fl-stop-fl^*;*Pten^fl/fl^* mice (coordinates: AP -1.0 mm and right -2.5 mm; depth 2.5-3 mm from the dura surface). For survival analyses, mice were monitored for signs of end-stage tumor burden such as >20% weight loss, visible bump on head, lethargy, and seizures. All mice were sacrificed at a humane endpoint with an overdose of ketamine (300 mg/kg) and xylazine (30 mg/kg) and perfused with Ringer’s solution.

### Human GBM samples

Formalin-fixed and paraffin-embedded (FFPE) human GBM samples and de-identified clinical information were provided by the Duke Cancer Institute (IRB #00114085). Sample information is provided in Supplementary Table 1. Edema scoring was done by a board-certified neuro-oncologist to determine extent of edema pre-operatively. Fresh-frozen tumor tissues used for Western blot analysis were collected at Mount Sinai Hospital as previously described (55) through the biorepository under protocols approved by the IRB (#18-00983). All patient samples were de-identified. The samples included in this study with molecular information are summarized in Supplementary Table 2.

### Wet/dry assay for measurement of edema content

End-stage, tumor-bearing mice were anesthetized with ketamine (300 mg/kg) and xylazine solution (30 mg/kg), whole brains were extracted without perfusion and placed on a pre-weighed piece of aluminum foil. The weight of the brain and foil was recorded as the wet weight. The brains were immediately placed into a vacuum oven (Across International) and allowed to dry at 95°C for 24 hours. After 24 hours, the weight of the brain and foil was recorded as dry weight. The percent brain water was calculated as [wet weight - dry weight)/dry weight]*100.

### MRI tumor volume reconstruction

MRI imaging was performed at the Imaging Core as part of the BioMedical Engineering and Imaging Institute (Icahn School of Medicine at Mount Sinai). All MRI data were acquired using a 9.4T Bruker scanner as previously described by Herting *et al*, 2019 (14). To determine tumor volume, we used Paravision 6.0 software and employed the freehand region tool to calculate the area of visible tumor burden on each slice. We then multiplied the area of each slice by the thickness of the slice (0.7 mm) to obtain the volume of tumor burden on the slice. To obtain the total tumor burden for the mouse, we summed all the resulting volumes from the scans.

### H&E reconstruction of tumor volume

End-stage tumor-bearing mice were sacrificed using an overdose of ketamine (300 mg/kg) and xylazine solution (30 mg/kg) and perfused transcardially with 4% paraformaldehyde (PFA) in PBS. Whole brains were collected in 4% PFA for 24 hrs followed by 72 hrs in 30% sucrose at 4°C. The brains were embedded in OCT (Fisher #23-730-571) on dry ice and serially sectioned on a Leica cryostat at 30 µm thickness. Every tenth section was collected on a slide. For H&E staining, slides from the freezer were immediately placed in cold 4% PFA solution for 10 minutes, followed by rinsing in 1X PBS and tap water to remove excess OCT. Slides were stained in Mayer’s Hematoxylin (Fisher #50-318-61) for 1 minute, rinsed with tap water, and placed in 1X PBS for 30 seconds to allow bluing of the stain. The slides were then passed through 70% and 95% ethanol solutions before placing them in alcoholic Eosin solution (Sigma #SDHT110132) for 30 seconds. Finally, slides were passed through several changes of 95% and 100% ethanol as well as Neo-Clear solution (Sigma #64741-65-7) before mounting using Permount medium (Fisher #SP15-100). The slides were scanned at 20X magnification with a whole slide scanner (Hamamatsu) and tumor area was outlined using NDP.View2 software. The tumor area of each section was multiplied by the thickness of 10 sections, and the resulting volumes were summed to calculate the total tumor volume for each sample.

### Measurements of intracranial pressure (ICP)

The setup involved a miniature pressure catheter (Millar Mikro-Tip Catheter Transducers, SPR 1000) consisting of a polyimide transducer tip that is suitable for pressure measurements in mice. This catheter was connected to a software-controlled bridge amplifier (AD Instruments #FE221) via a Mikro-Tip Catheter Interface Cable. The bridge amplifier allowed for pressure readings from the transducer to be visualized on a computer screen using the LabChart software. To measure ICP, mice at 4-6 weeks of age were injected with a combination of plasmids to generate *de novo* PDGFB mGBM. Age-matched naive mice were used as controls. When the mice developed endpoint symptoms such as >20% weight loss, a visible bump on the head, and hunched appearance, they were anesthetized using ketamine (100 mg/kg) and xylazine (10 mg/kg). A hole was drilled in the right frontal striatum at the site of injection, followed by lowering the tip of the probe into the brain at a depth of 1.5 mm. Intracranial pressure was recorded for approximately 10 minutes per mouse, allowing the first few minutes for probe stabilization. The mice were sacrificed after measurement, and the brains were banked in formalin without perfusion.

### Immunohistochemistry staining

Brains were collected from tumor-bearing mice post-perfusion and placed in 10% formalin solution for 72 hours, followed by 1X PBS until paraffin-embedding. The paraffin blocks were cut as 5 µm sections on a microtome (Leica) and collected directly on SuperFrost glass slides (Fisher #22037246). For each set of stainings, antibodies at the pre-standardized dilutions (Supplementary Table 4) and an automated stainer (Leica, Bond RX) were used. Post-staining, the slides were dehydrated by passing through increasing grades of alcohol and Neo-Clear followed by mounting in Permount medium. Mounted slides were allowed to dry overnight and scanned at 20X magnification with a whole slide scanner (Hamamatsu). NDP.View2 software was used for capturing multiple screenshots representative of the tumor and quantification was done using the ImageJ software.

### Immunofluorescence staining and confocal microscopy

For formalin-fixed paraffin-embedded (FFPE) samples, the slides were deparaffinized by passing through Neo-Clear and decreasing grades of ethanol followed by antigen retrieval in a boiling solution of citrate-based buffer at pH 6 (Vector Labs #H-3300). The sections were allowed to cool on ice and then subjected to permeabilization with 0.3% Triton-X 100 detergent solution in PBS. Blocking was performed for 1 hour at room temperature (RT; or overnight at 4°C) using 0.1% Triton-X 100 containing 2% BSA and 5% normal donkey serum (Jackson ImmunoResearch). The primary antibodies were diluted in PBS (Supplementary Table 4) and incubated for 1 hour at RT or overnight at 4°C. Secondary antibodies (Jackson ImmunoResearch) were used at 1:500 dilution in PBS for 1 hour at RT, followed by counterstaining with DAPI and mounting in 70% glycerol solution. All images were taken on the Leica SP8 confocal microscope, keeping laser and software settings consistent across samples. Images were analysed using the ImageJ (FIJI) software.

### Orion multiplex IF staining

FFPE tissue sections were deparaffinized, rehydrated, and subjected to antigen retrieval by submersion in retrieval buffer (BioGenex #HK547-XAK) followed by microwaving (BioGenex, EZ-RT-MW016-IR) for 10 minutes. Non-specific protein binding was blocked using a mixture of normal sera derived from mouse (Millipore Sigma #M5905), rabbit (Millipore Sigma #R4505), and goat (Fisher #16210064). A cocktail containing 13 fluorophore-conjugated antibodies (Supplementary Table 6) was then applied to the slides and incubated for 2 hours at room temperature. Unbound antibodies were removed by washing. Hoechst 33342 (Fisher #H3570) was added to the sections at 1:500 dilution for 30 min at RT. Slides were cover slipped with ArgoFluor™ Mounting Medium (RareCyte #42-1214-000) and loaded onto the Orion platform (100) for image acquisition. Whole-slide imaging was performed on the Orion (RareCyte) system equipped with seven high-power excitation lasers and sensitive CMOS detectors, coupled with optical filters providing 90–95% transmission efficiency. These filters enabled the collection of 10–15 nm bandpass channels with 1-nm center wavelength (CWL) tuning selectivity. Following image acquisition, tiles were stitched, distortion-corrected, and processed using a pre-established spectral unmixing matrix. Matrix coefficients were derived by scanning single-stained spectral controls to capture both target fluorophore emission and off-target fluorescence across all Orion channels. Using RareCyte’s Artemis software, proprietary algorithms calculated correction factors for spectral crosstalk between donor and recipient fluorophores, as well as for tissue and background autofluorescence. The finalized spectral unmixing matrix was then applied to multiplexed image data to retain true fluorophore-specific signal intensity while removing non-specific crosstalk and autofluorescence. Image files were exported as pyramidal, multi-layered OME-TIFF files.

### Orion multiplex IF data analysis

Fluorescence image segmentation was performed using the StarDist Python API (v0.9.1) in Python v3.10.18, applied to the Hoechst nuclear channel. Segmented nuclei were subsequently expanded by up to 5 µm from the nuclear boundary to approximate cytoplasmic regions, resulting in whole-cell masks. Following segmentation, cell masks were overlaid with raw fluorescence images, and the mean pixel intensity per cell was computed for each channel. Morphometric features -- including nuclear area, eccentricity, solidity, and centroid coordinates (X, Y) -- were extracted from the nuclei channel using the scikit-image Python package (v0.25.2) (101). Channel-specific technical artifacts were visually inspected and removed using napari (Version v0.6.6rc2). Segmented cell masks were overlaid with log-normalized fluorescence signals to determine channel-specific thresholds distinguishing positive and negative cells. These thresholds were subsequently used to rescale fluorescence intensities to a 0-1 range using the scimap Python package (v2.3.5) (102). Cells with scaled channel values ≥ 0.5 were classified as positive for that marker. Composite marker-positive regions were defined using Boolean logic operations. For example, the IBA-positive area was defined as region, where (IBA1≥0.5) and (GFAP<0.5) and (ALDH1L1<0.5) and (OLIG2<0.5) and (CD8a<0.5) and (CD31<0.5) and (CD15<0.5) and (CD20<0.5). Applying exclusion thresholds of < 0.5 for non-target markers minimized false-positive assignments arising from overlapping or nonspecific signals.

### Western Blot

RIPA lysis buffer containing protease-inhibitor cocktail was added to snap-frozen tissue or frozen cell pellets (Supplementary Tables 2&3) followed by sonication until the tissue was completely homogenized. The samples were incubated on ice for 30 minutes and then centrifuged at 13,000 rpm for 10 minutes at 4°C. The supernatant was collected in a fresh tube and stored at -80°C until required. To determine protein concentration, the Bradford assay was used. 30 μg of denatured protein and a pre-stained protein ladder was loaded on a pre-cast 4-12% SDS-PAGE gel (Invitrogen #NW04120BOX) and run at 200 V for approximately 30-40 minutes depending upon the separation required. The separated proteins were then transferred to a PVDF membrane under cold conditions for 1 hour at 100 V. The membrane was blocked for 1 hour at RT using 5% milk in TBST (0.1%; CST #9997) and the following primary antibodies were incubated for 1 hour at RT on a shaker: anti-AQP4 (1:10,000; Proteintech #16473-1-AP) and anti-β actin (1:50,000; CST #3700S). Following three 10-min washes in TBST, the membrane was incubated with species-specific secondary antibodies (1:100,000; CST #7074 & #7076) for 1 hour at RT on a shaker. The membrane was washed three times with TBST and incubated with ECL (Fisher #34075) for 5 minutes and developed in a dark facility on an X-ray film (Fisher #PI34090).

### CD4^+^ and CD8^+^ T cell depletion

For the CD8 depletion cohort, 200 μg of anti-CD8 antibody (Clone 2.43, BioXCell #BE0061) or IgG2b (BioXCell #BE0090) control prepared in sterile 1X PBS (Fisher, #J62036.K2) was administered every 4 days starting at day 20 post-injection and continuing until endpoint signs developed. CD4 depletion was performed as previously described (73). Briefly, 300 µg of anti-CD4 antibody (Clone GK1.5, BioXCell #BE0003-1) or IgG2b control (BioXCell #BE0090) prepared in sterile 1X PBS (Fisher, #J62036.K2) was initially administered at day 22 post-injection and again two days later, followed by treatment every 7 days until endpoint signs developed.

### Spectral flow cytometry

Brains from tumor-bearing mice were collected on ice post-perfusion with ice cold Ringer’s solution and the tumor was dissociated using 0.5% collagenase D and DNase I. The single cell suspension was subjected to density-gradient centrifugation with 30% Percoll solution containing 10% FBS. After careful aspiration of the supernatant, the pellet was resuspended in RBC Lysis Buffer for 1 minute, centrifuged, and the resulting pellet was resuspended in 100 µl of blocking buffer (2% FBS, 5% normal rat serum, 5% normal mouse serum, 5% normal rabbit serum, 10 μg/ml anti-FcR (BioLegend #101319) in FACS Buffer). Blocking was performed for 20 minutes on ice, and the suspension was split into two tubes before centrifugation. The supernatant was aspirated from each pellet, and one tube was incubated with primary antibodies from the myeloid panel (Supplementary Table 5) while the other was incubated with antibodies from the lymphoid panel (Supplementary Table 5). Primary antibody incubation was carried out for 30 mins on ice, followed by a wash with PBS-EDTA solution and then resuspended in 100 µl of viability dye (Zombie UV; Biolegend #423108) prepared in PBS-EDTA for 20 mins at RT. Cells were then washed with FACS buffer (0.1% BSA and 0.1% sodium azide in 1X PBS) and fixed with fixation buffer (eBioscience #5223-56, #5123-43) for 30 mins at 4°C. The cells were washed again and incubated with a mixture containing intracellular antibodies (Supplementary Table 5) in permeabilization buffer (Invitrogen #8333-56) for 30 mins at RT. Finally, cells were washed, resuspended in 300 µl of FACS buffer and stored at 4°C until analysis. The samples were run on an Aurora Cytek spectral flow cytometer, and all data were analyzed using FlowJo software (Tree Star Inc, Version 10). For evaluating levels of circulating immune cells, blood was extracted from anesthetized mice retro-orbitally and 50 µl was combined with an equal volume of antibodies (Supplementary Table 5). The solution was incubated for 15 minutes at RT, followed by RBC lysis for 10 minutes at RT. Counting beads (Invitrogen #C36995; 25 µl) were added and the samples were run on an Aurora Cytek cytometer on the same day.

### Flow cytometry for GFP-positive astrocytes from *Aldh1l-eGFP* mice

Brains from naive or end-stage tumor-bearing *Ntv-a; Aldh1l1-eGFP*^+/+^ mice were collected on ice post-perfusion with ice-cold Ringer’s solution. Cortex or tumor tissue, respectively, was minced into small pieces and subjected to enzymatic digestion with a mixture of papain (0.94 mg/ml, Worthington #LS003120), 0.5 M EDTA, DNase (60 ug/ml, Roche) and 0.5 M N-Acetyl-cysteine prepared in EBSS solution. The tissue was digested for 15 minutes at 37°C followed by termination of the process using Ovomucoid solution (0.7 mg/ml, Worthington #LS003086). The suspension was pelleted by two rounds of centrifugation at 450 rcf, 5 mins and 600 rpm, 5 mins, respectively, to remove excess debris. A single cell suspension was obtained by passing the solution through a 70 µm cell strainer and into a 30% Percoll gradient followed by centrifugation to remove myelin debris. The resulting pellet was resuspended in 100 µl of blocking buffer and incubated on ice for 20 minutes. After blocking, the pellet was incubated with antibodies against CD45 and CD11b for 30 minutes on ice. Viability dye staining (Zombie UV; Biolegend #423108) was performed at RT for 20 minutes followed by a wash with FACS buffer. The pellet was finally resuspended in 300 µl of FACS buffer and run immediately on the Aurora Cytek spectral flow cytometer. All data were analyzed using the FlowJo software.

### RNAscope fluorescent *in situ* hybridization and co-detection with immunohistochemistry

This experiment was performed in collaboration with Dr. Jia Liu at the CUNY Advanced Science Research Center. FFPE blocks of tumor-bearing murine brain tissue were serially cut using a microtome (5 μm thickness) on SuperFrost slides and kept at RT until further processing. *In situ* hybridization combined with immunohistochemistry was performed using RNAscope probes and reagents from Advanced Cell Diagnostics (ACD) according to the manufacturer’s instructions. Briefly, sections were first deparaffinized using xylene and ethanol. Endogenous peroxidase activity was quenched with hydrogen peroxide reagent for 10 min, followed by antigen retrieval for 10 min in boiling buffer. Immunohistochemistry was performed afterwards using anti-OLIG2 (Rabbit Millipore #AB9610, 1:100) and anti-GFAP antibodies (Mouse CST #3670T, 1:250), followed by protease digestion for 30 min at 40°C. The following probe was then hybridized for 2 h at 40°C in a humidity-controlled oven (HybEZ II, ACD): RnAqp4 (C2). Successive addition of amplifiers was performed using the proprietary AMP reagents, and the signal visualized through probe-specific horseradish-peroxidase-based detection by signal amplification with Opal dye 570 (Perkin Elmer, 1:1500). Sections were incubated with Donkey anti-Mouse (Alexa Fluor 488, 1:500) and Donkey anti-Rabbit (Alexa Fluor 647, 1:500) secondary antibodies at RT. Slides were then counterstained with DAPI and mounted with Prolong Gold Antifade (Fisher #P36930) and kept at 4°C until imaging. Slides were imaged on a Leica Stellaris 5 confocal microscope keeping laser and software settings consistent between samples.

### Vessel leakage assay

B20-4.1.1 (Genentech) antibody was administered to tumor-bearing mice intraperitoneally (5 mg/kg) at symptom onset and was continued every 4 days until endpoint. At endpoint, the mice were anesthetized with Ketamine (300 mg/kg) and Xylazine solution (30 mg/kg) and given retro-orbital injections of FITC-Lectin (50 µl, Vector Labs #DL-1174) for 5 mins and Hoechst dye (50 ul, Sigma #94403) for an additional 5 mins. The mice were then transcardially perfused with ice-cold Ringer’s solution followed by 4% PFA solution. The brains were collected in 4% PFA and fixed for 24 hours, after which they were transferred to 30% sucrose solution for 72 hours before embedding in OCT. Sections (8 µm) were cut on a cryostat (Leica) and collected directly on SuperFrost slides. The sections were permeabilized by incubating with 0.3% Triton X-100 for 30 minutes at RT, followed by three washes with 1X PBS. For propidium iodide (PI) staining, a mixture of PI (Fisher #BDB556463), RNase, and Triton X-100 in PBS was prepared and the slides were incubated in this solution for 30 minutes at 37°C. After washing three times with 1X PBS, the slides were mounted in 70% glycerol and stored at 4°C until imaging. All imaging was performed on a Leica SP8 confocal microscope, keeping laser and software settings consistent across samples. Image analysis was performed using FIJI software. Briefly, the tumor area was outlined using the PI staining as a guide, and the Hoechst-positive areas were individually outlined and quantified using the freehand region tool. To calculate the percentage of Hoechst-positive area within the tumor, the individual area values for each sample were summed and normalized to the tumor area for that sample.

### Quantitative PCR

RNA was extracted from a snap-frozen tumor or naive brain tissue using Qiazol (Qiagen #79306) and RNeasy kit (Qiagen #74104) according to the manufacturer’s instructions. The extracted RNA was quantified using a NanoDrop 2000 spectrometer and run on a 1.5% agarose gel containing 1% bleach to assess quality. 1 µg of RNA was used for cDNA synthesis using the First Strand Superscript III cDNA synthesis kit (Invitrogen #18080051) according to manufacturer’s instructions. Quantitative PCR was run for the cDNA samples using primers targeting the *Aqp4* gene (BioRad #10025636) and SsoAdvanced Universal Green Supermix (BioRad #1725272). Gene expression was quantified relative to the expression of housekeeping gene *β-actin* followed by fold change calculation using the 2^-ΔΔCt^ method.

### Statistical Analysis of MRI Features by Molecular Subtype

To evaluate associations between molecular subtypes and *AQP4* expression, as well as MRI-derived imaging features, we conducted statistical analyses in R (v4.3.0 or later) using pre-processed and batch-corrected bulk RNA-seq data (n = 159) from multiregional human GBM samples (36). *AQP4* expression values were z-score-normalized across all samples. Patient-level metadata, including transcriptionally-defined Verhaak subtype annotations and imaging features (e.g., T2-weighted signal intensity), were used in downstream analysis. Samples labeled as “not classified” (nc) under the Verhaak classification were excluded. Pearson correlation tests were used to assess the relationship between T2 signal and *AQP4* expression across all samples, as well as within the contrast-enhancing (CE) subgroup. Linear mixed-effects models were applied to evaluate the effect of Verhaak subtype (Classical, Mesenchymal, Proneural) on *AQP4* expression and T2-weighted MRI intensity (T2.Raw_Mean), using the lmer() function from the lme4 package (v1.1-37), with Patient ID included as a random intercept to account for repeated measures. Fixed effects were visualized using model estimates and 95% confidence intervals (CIs), displayed as point-range plots. P-values were reported to indicate significance while controlling for inter-patient variability. Type II ANOVA was performed using the lmerTest package (v3.1-3), and pairwise *post hoc* comparisons between subtypes were computed using the emmeans package (v1.11.0). Visualization of results was carried out using ggplot2 and ggpubr, with global ANOVA p-values and pairwise contrast p-values annotated on relevant plots.

### Single-cell RNA sequencing

Tumor-bearing *Ntv-a;Aqp4^+/+^* and *Ntv-a;Aqp4^-/-^*mice at endpoint were perfused with ice cold Ringer’s solution and the brain was immediately collected on ice. The tumor was minced into smaller pieces and subjected to enzymatic digestion using papain/ovomucoid resulting in a single cell suspension as described above. Cell viability was determined using Trypan Blue and samples with greater than 80% viability were selected for sequencing. All sequencing was performed at the Genomics Core (ISMMS) using the 3’ GEX V3 kit and Chromium platform (10X Genomics). Raw 10X Genomics data were processed with the vendor pipeline to generate gene–barcode matrices (Cell Ranger). Count matrices were imported and combined into a unified object, preserving batch and condition metadata (Seurat and scanpy) (103, 104). Quality control used per-cell metrics (library size, detected features, mitochondrial proportion) with visual inspection to define conservative thresholds; subsequent analyses used only high-quality cells. Library-size normalization relied on counts-per-cell scaling followed by log transformation to stabilize variance and approximate homoscedasticity. Highly variable genes were selected to capture biological signal over technical noise, data were centered and scaled per gene, and dimensionality was reduced by principal component analysis. To correct inter-batch effects while preserving biological structure, we performed integration by removing shared linear factors of unwanted variation in the low-dimensional space (Harmony) (105). Graph construction used a k-nearest neighbor graph on the integrated embedding, and communities were identified by graph-based modularity optimization. Nonlinear manifold learning was applied for visualization by Uniform Manifold Approximation and Projection, which preserves local neighborhood structure. Cell-type annotation combined supervised reference mapping and de novo curation. First, query cells were projected onto a curated brain reference using anchor-based transfer to obtain predicted labels at multiple hierarchical levels (Azimuth) (106). Second, cluster-level marker discovery employed a nonparametric rank-sum test with multiple-testing correction to identify discriminative genes. Module scores were computed for predefined astrocyte programs to derive per-cell signature activity. For focused analyses, astrocytes were subset, re-embedded, reclustered at low resolution, and contrasted between conditions. In parallel, we incorporated a probabilistic deep generative embedding that models gene counts with a negative binomial likelihood and amortized inference to capture nonlinear biological variation (scvi-tools) (107). The learned low-dimensional representation was imported, used for neighbor graph construction, clustering, and UMAP, and contrasted with the Harmony workflow to confirm stability of cell states across embeddings. Pathway analysis used preranked gene set enrichment: genes were ordered by signed effect size from differential testing, and enrichment was assessed against curated ontologies to identify coordinated biological programs with permutation-based statistics. Cell–cell communication was inferred from ligand–receptor expression using a probabilistic framework that computes interaction probabilities per pair of cell groups, aggregates signals at pathway level, and quantifies network centrality to identify sending/receiving hubs; similarities between signaling programs were embedded and clustered for comparative analysis (CellChat) (108).

### Spatial transcriptomics

Tumor-bearing *Ntv-a;Aqp4^+/+^* and *Ntv-a;Aqp4^-/-^*mice at endpoint were perfused with ice cold Ringer’s solution and the brain was immediately collected in 10% formalin for 6 hours at RT. The tissue was then transferred to 1X PBS and stored in the fridge for 24-48 hours. The samples were then embedded in paraffin. FFPE tissue blocks were sectioned for both RNA quality assessment and spatial transcriptomics. For RNA extraction, five 10 µm curls were deparaffinized, digested with proteinase K, treated with DNase, and purified by silica-membrane spin columns (RNeasy FFPE, Qiagen #73504). RNA integrity was evaluated using capillary electrophoresis, and the DV200, representing the fraction of RNA fragments between 200 and 10,000 bp, was calculated (ProSize) to confirm suitability for spatial analysis (DV200 > 50%). For spatial transcriptomics, 5 µm FFPE sections were rehydrated, floated on a 42 °C water bath, mounted onto Poly-Prep slides within the capture area, and baked at 42 °C for 3 h before desiccated storage. Prior to assay processing, slides were heated to 60 °C, deparaffinized in xylene, and rehydrated through graded ethanol series. Sections were briefly hematoxylin- and eosin-stained, imaged by brightfield microscopy (10X), and then destained for subsequent molecular processing. Probe hybridization and library preparation followed the Visium CytAssist FFPE protocol (10X Genomics). After acid destaining and heat-induced decrosslinking, species-specific probe sets were hybridized to the tissue overnight at 50 °C, ligated, and washed under stringent thermal conditions. Released probe pairs were extended, eluted under alkaline conditions, neutralized, and amplified with pre-amplification and indexing reactions. Libraries were size-verified and quality-controlled by electrophoresis (TapeStation) prior to sequencing. This workflow ensured optimal RNA recovery, spatial RNA capture efficiency, and robust library construction from archival FFPE material.

### Spatial transcriptomics Data Analysis

Spatial transcriptomics datasets (Visium) were processed in a unified framework to enable cross-modality comparisons despite resolution differences. Count matrices were imported into a spatial object model that preserves per-spot coordinates, histology alignment, and per-spot cell-type estimates from single-cell reference deconvolution (SPATA2; Cell2location; Cytospace) (81, 82, 109). Mouse–human gene symbol harmonization was performed where required using curated orthology tables prior to downstream analyses. Expression normalization used a probabilistic latent variable model that accounts for library size and over-dispersion, yielding denoised gene abundances and a nonlinear low-dimensional embedding of biological variation (negative-binomial variational autoencoder; scVI via scvi-tools) (107). The normalized matrix was set as the active expression layer for all spatial summaries and gene-set scoring. Tissue compartmentalization was derived by rule-based spatial segmentation integrating image layers and transcript features to assign tumor versus non-tumor regions on the native grid (SPATA2). Spatial expression maps for genes and modules were rendered as continuous fields with value clipping to reduce outlier influence. For ecosystem discovery, we constructed spatial graphs on each section by connecting spots within a fixed Euclidean distance threshold, then extracted multi-hop (3-hop) subgraphs to capture local neighborhoods. Nodes encoded normalized gene vectors; edges carried inter-spot distances. A multi-head graph attention encoder learned subgraph representations with edge-aware attention, followed by global mean pooling to obtain a single embedding per subgraph as recently published by our group (83). A multitask head jointly (i) classified genotype status (*Ntv-a;Aqp4^+/+^* and *Ntv-a;Aqp4^-/-^*) and (ii) regressed a continuous region score reflecting tumor proximity, optimizing a composite loss that combined cross-entropy and a robust regression objective (Huber) with early stopping to prevent overfitting. Model performance was quantified by accuracy, F1, confusion matrices, and ROC–AUC on held-out subgraphs. To interpret predicted ecosystems at macroscopic scale, subgraph scores were mapped back to the tissue and visualized over histology to identify high-confidence regions enriched for KO-like states. For biological characterization, we generated pseudobulk profiles by summing counts over all spots belonging to each ecosystem, projected these into a common gene space, and performed differential expression between *Ntv-a;Aqp4^+/+^*and *Ntv-a;Aqp4^-/-^* enriched ecosystems using a non-parametric rank-based test with multiple-testing correction. Preranked enrichment analysis then identified coordinated pathways across ontologies using signed effect sizes (GSEA to GO/KEGG; clusterProfiler/DOSE). Cell state programs were quantified by module scoring of predefined astrocyte signatures to position astrocytic neighborhoods along activation and maturation axes. Local cell–cell milieu was examined by nearest-neighbor analysis around astrocyte seeds to summarize contact frequencies with other lineages, followed by weighted network visualization to highlight genotype-specific shifts in neighborhood composition. All spatial figures used consistent axes with fixed aspect ratios; color scales were chosen to preserve perceptual ordering and comparability across samples.

### Statistical analysis

All statistical analyses were performed using GraphPad Prism software (Version 9). P-values less than 0.05 were considered to be statistically significant. For details relating to specific experiments, please refer to the figure legends.

## Data availability

Genomic data including the spatial transcriptomics data from the Visium platform and the single cell RNA-sequencing data are available 10.5281/zenodo.17597432. Code implementation for analysis is available at github (https://github.com/heilandd/AQP4_Project).

## Acknowledgements

We thank all members of the Hambardzumyan laboratory for technical assistance and constructive discussion of the results. We would also like to acknowledge the Microscopy and Advanced Bioimaging Core, Flow Cytometry Core, and the Genomics Core Facilities at the Icahn School of Medicine at Mount Sinai for providing resources and assistance for confocal imaging, spectral flow cytometry, and single cell sequencing. P30 CA196521 supports the Tisch Cancer Institute and related research facilities. We thank Yu Zhou at the Biomedical Engineering and Imaging Institute (ISMMS) for assistance with MRI set-up, and Dr. Jia Liu at the CUNY Epigenetics Core Facility for assistance with the RNAscope experiment. This work was supported by NIH/NINDS (R01NS100864) funding to DH. DHH is supported by the Heisenberg Program of the DFG (HE 8145/5-1 & HE 8145/5-2), Cancer Consortium (DKTK) and the Transcan PLASTIC project (D.H.H.), funded by BMBF (iGerman Ministry of Education and Research). Graphical abstracts in all figures were created using BioRender. We appreciate Chris Nelson for his editing contributions.

## Author Contributions

The indicated authors were responsible for the following roles. Designing research studies: all authors. Conducting experiments: T.J., K.G., M.J.F., M.P.V., R.R., A.J.S., D.H. Analyzing data: T.J., M.J.F., K.G., M.D., S.I.R., E.R., D.D., K.B.P., and D.H.H. Providing intellectual input, samples, and reagents: Z.C., K.B.P, N.M.T., N.L.T., L.S.H., A.J.S., D.H.H., and D.H. Writing the manuscript: T.J., D.H.H., and D.H. Editing the manuscript: all authors. T.J. was responsible for creating all the figures with input from M.J.F., D.H.H and D.H. D.H. conceptualized and supervised the study and obtained funding.

## Extended Figures

**Extended Data Fig. 1.**
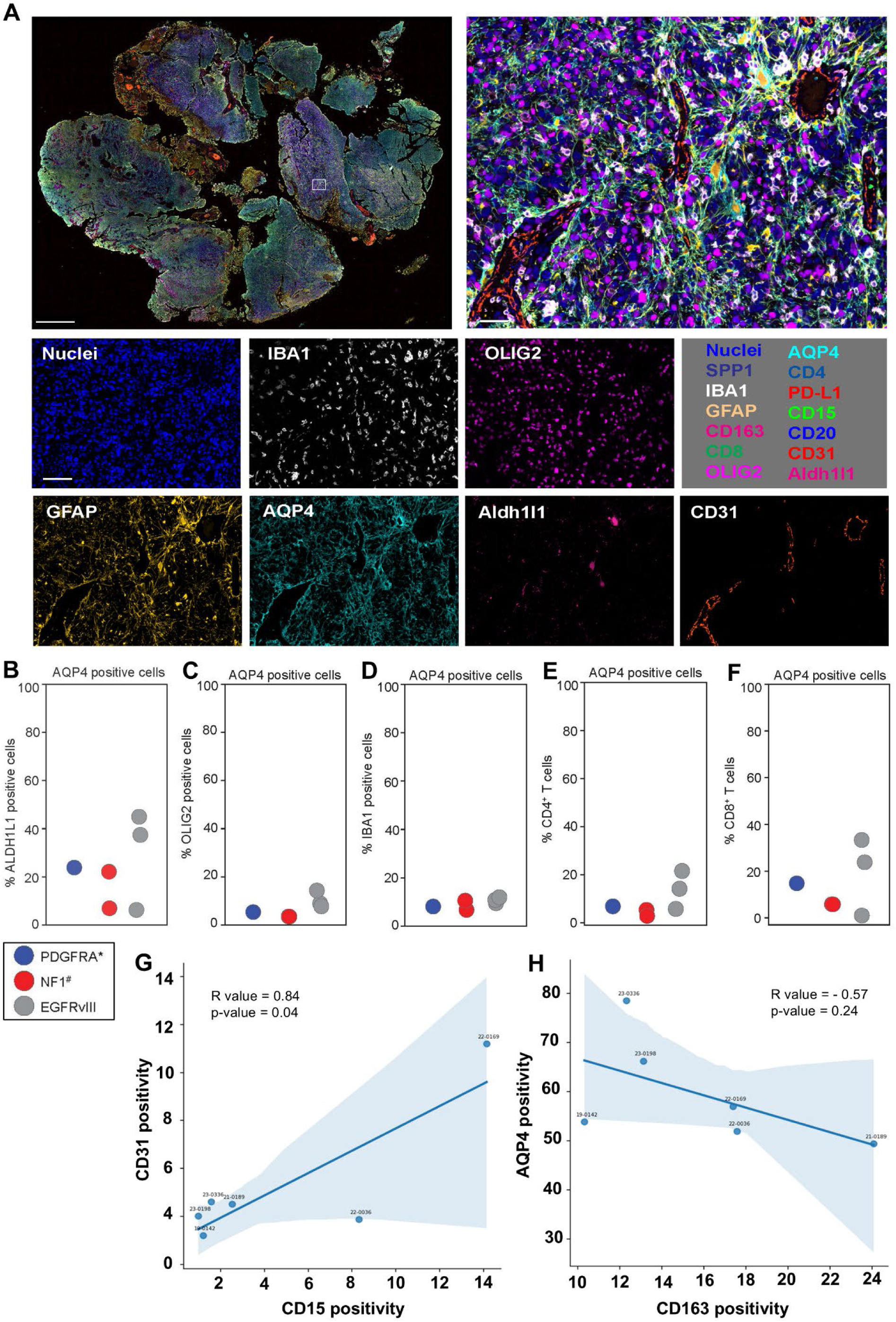
Multiplex IF staining of human GBM samples using the Orion^™^ platform. **A**, Representative whole mount scan and high magnification insets of multiplex IF staining on a human GBM tissue section for various markers using the Orion platform. Scale bar = 2 mm, 50 μm, 160 μm (left to right and top to bottom). **B-F**, Graphs showing AQP4 positive cells as a percentage of ALDH1L1-, OLIG2- , IBA1-, CD4-, and CD8-positive cells respectively, quantified by Orion multiplex IF staining in human GBM samples (n=6). Legend indicates whether the samples contain *PDGFRA* amplification, *NF1* mutation or *EGFRvIII* mutation. **G,** Linear regression analysis of CD31 and CD15 positivity in human GBM samples (n=6). Each point represents an individual sample. The solid line represents the linear regression fit. Pearson’s correlation analysis (R value = 0.84, P = 0.04). **H**, Linear regression analysis of AQP4 and CD163 positivity in human GBM samples (n=6). Each point represents an individual sample. The solid line represents the linear regression fit. Pearson’s correlation analysis (R value = -0.57, P = 0.24).

