## Supplementary Figures and Tables for "Aquaporin-4 Regulates Edema, Sustains Astrocyte Differentiation, and Constrains Immunosuppressive Remodeling in Glioblastoma"

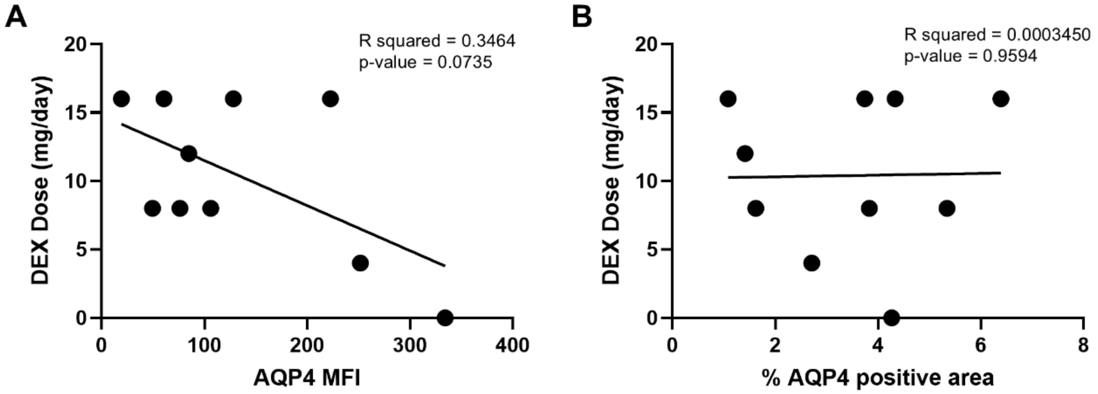


C

| Sample ID | Edema Score | EGFR VIII | PDGFRA Amplification | NF1 | TERT |
|---|---|---|---|---|---|
| 21-0189 | Mild | Variant Not Detected | No Amplification | Mutated, Pathogenic | Mutation Present |
| 19-0051 | Mild | Variant Detected | No Amplification | Indeterminate | Indeterminate |
| 19-0142 | Mild | Variant Detected | No Amplification | Indeterminate | Indeterminate |
| 23-0170 | Moderate | Variant Detected | No Amplification | No Mutation | Mutation Present |
| 22-0036 | Severe | Variant Detected | No Amplification | No Mutation | Indeterminate |
| 22-0169 | Mild | Variant Not Detected | No Amplification | Mutated, Pathogenic | Indeterminate |
| 23-0198 | Severe | Variant Detected | No Amplification, But Variant of Uncertain Significance | No Mutation | Mutation Present |
| 21-0057 | Moderate | Variant Detected | No Amplification | No Mutation | Mutation Present |
| 23-0093 | Moderate | Variant Detected | No Amplification | No Mutation | Mutation Present |
| 23-0336 | Mild | Variant Not Detected | Amplification Present | No Mutation | Mutation Present |
| A15-88 | - | No Mutation | No Mutation | No Mutation | No Mutation |
| A15-014 | - | No Mutation | No Mutation | No Mutation | No Mutation |
| A15-107 | - | No Mutation | No Mutation | No Mutation | No Mutation |
| A15-109 | - | No Mutation | No Mutation | No Mutation | No Mutation |

**Supplementary Fig. 1. Linear regression analysis of AQP4 and dexamethasone dose in GBM patients. A**, Scatter plot depicting the association between DEX dose (mg/day) and AQP4 mean fluorescent intensity (MFI) determined by IF staining in human GBM samples. Each point represents an individual sample. The solid line represents the linear regression fit. Pearson's correlation analysis (R squared = 0.3464, P = 0.0735). **B**, Scatter plot depicting the association between DEX dose (mg/day) and AQP4 positive area (percentage) determined by IF staining in human GBM samples. Each point represents an individual sample. The solid line represents the linear regression fit. Pearson's correlation analysis (R squared = 0.0003450, P = 0.9594). **C,** Table with information on edema score and mutational profile of human samples used for correlation studies between DEX dose and AQP4 expression.

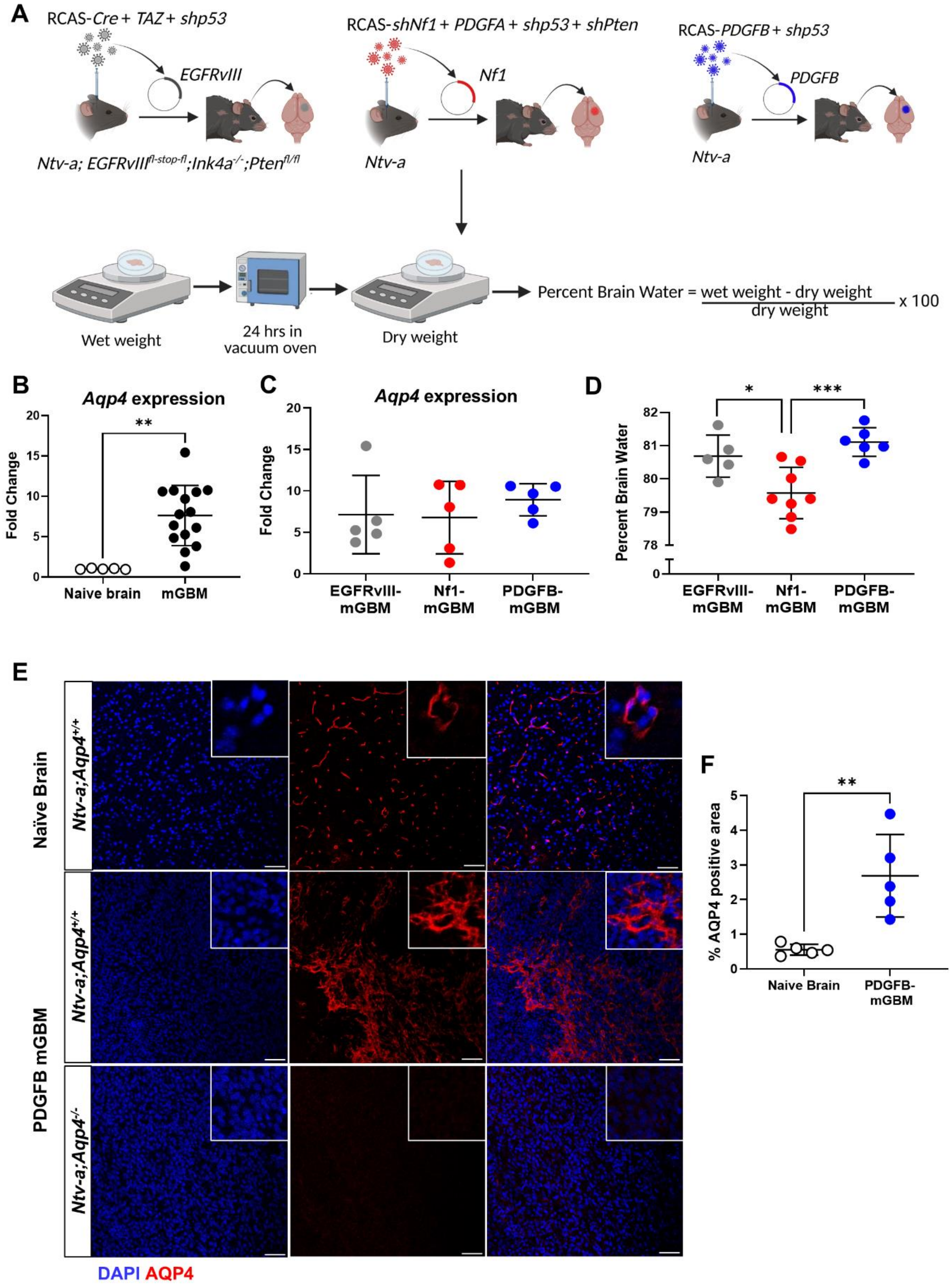


**Supplementary Fig.2. Evaluation of AQP4 expression in murine GBM. A**, Schematic illustration of the generation of murine GBM with different driver mutations using the RCAS/*tv-a* system, followed by edema measurement using the wet/dry assay. **B**, Graph showing fold change in *Aqp4* gene expression over housekeeping gene *β-actin*, compared between naïve murine brain cortex tissue and murine GBM tissue (mGBM). Two-tailed Student's t-test, $^{**}p<0.01$. **C**, Graph showing fold change in *Aqp4* gene expression over housekeeping gene *β-actin,* compared between EGFRvIII-,Nf1- and PDGFB-mGBM. Ordinary one-way ANOVA with multiple comparisons. **D**, Comparison of edema content as percent brain water between EGFRvIII-,Nf1- and PDGFB-mGBM. Ordinary one-way ANOVA with multiple comparisons, $^{*}p<0.05$, $^{***}p<0.001$. **E,** Representative immunofluorecence images showing AQP4 (red) protein expression in naïve murine brain sections and PDGFB mGBM-bearing *Ntv-a;Aqp4*$^{+/+}$ and *Ntv-a;Aqp4*$^{-/-}$ murine brain sections. DAPI (blue) was used as a nuclear stain. Images were taken on the Leica SP8 confocal microscope. Scale bar=100 um. **F**, Quantification of AQP4 staining as shown in E, compared between naïve mice and PDGFB mGBM-bearing *Ntv-a;Aqp4*$^{+/+}$ mice. Two-tailed Student's t-test, $^{**}p<0.01$.

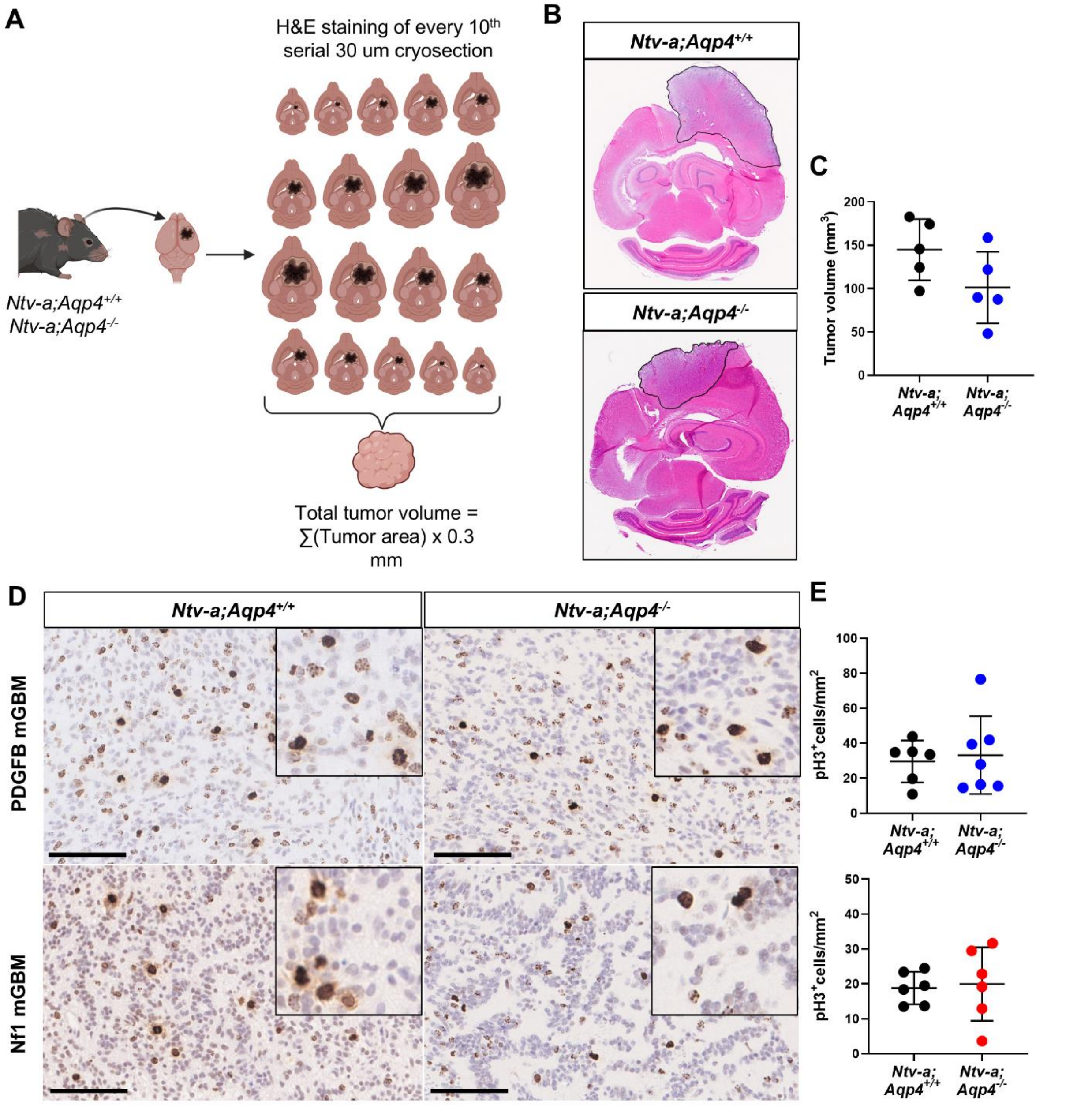


**Supplementary Fig.3.** ***Aqp4*** **loss affects tumor volume but not proliferation in murine GBM.** **A,** Schematic illustration of tumor volume reconstruction of MRI-imaged *Ntv-a;Aqp4*$^{+/+}$ and *Ntv-a;Aqp4*$^{-/-}$ mice bearing PDGFB mGBM using serial cryo-sectioning followed by H&E staining**.** **B**, Representative scans of H&E staining in MRI-imaged *Ntv-a;Aqp4*$^{+/+}$ and *Ntv-a;Aqp4*$^{-/-}$ mice bearing PDGFB mGBM. Slides were scanned at 20X magnification using a Hammamatsu slide scanner. **C,** Quantification of tumor volume in $mm^3$ as compared between MRI-imaged *Ntv-a;Aqp4*$^{+/+}$ and *Ntv-a;Aqp4*$^{-/-}$ mice bearing PDGFB mGBM . Two-tailed Student's t-test. **D,** Representative immunohistochemistry images of pHH3 staining in *Ntv-a;Aqp4*$^{+/+}$ and *Ntv-a;Aqp4*$^{-/-}$ mice bearing PDGFB- and Nf1 mGBM. Slides were scanned using a Hammamatsu slide scanner at 20X magnification, with insets at 40X magnification. Scale bar = 100 um. **E**, Graphs showing number of pHH3 cells per $mm^2$ compared between *Ntv-a;Aqp4*$^{+/+}$ and *Ntv-a;Aqp4*$^{-/-}$ mice bearing PDGFB- and Nf1 mGBM. Two-tailed Student's t-test.

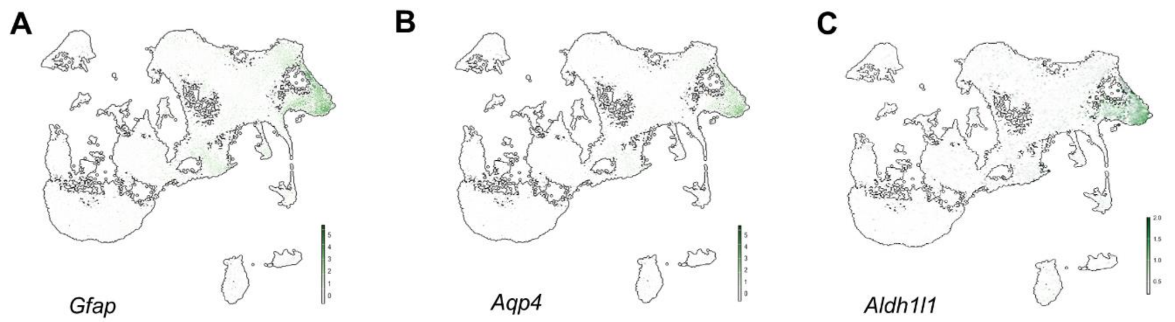


**Supplementary Fig.4. *Aqp4* co-localizes primarily with *Gfap* and *Aldh1l1* in murine GBM scRNAseq data. A-C,** UMAP dimensionality reduction plot showing expression of *Gfap, Aqp4* and *Aldh1l1* across all subclusters identified by scRNAseq analysis of n=4 *Ntv-a;Aqp4*$^{+/+}$ and n=3 *Ntv-a;Aqp4*$^{-/-}$ PDGFB mGBM tumors.

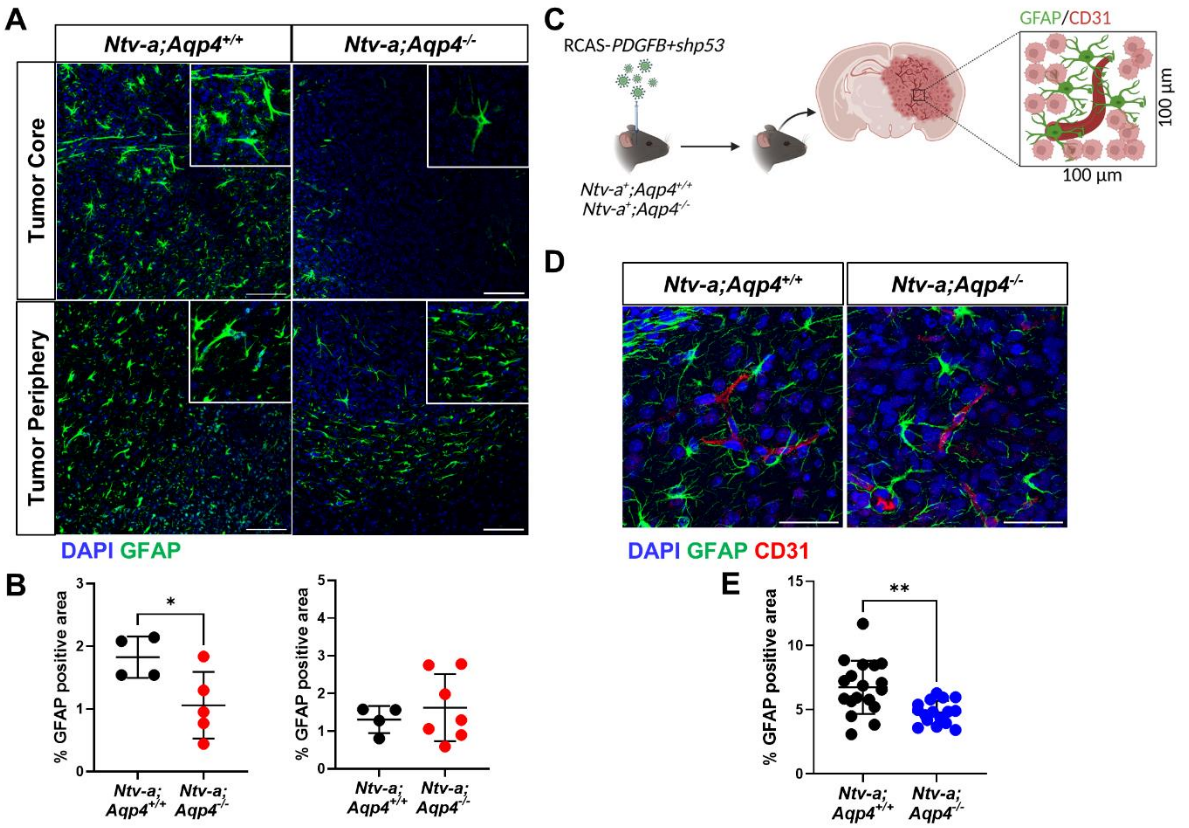


**Supplementary Fig.5.** ***Aqp4*** **loss results in decreased astrocyte coverage within the tumor core and in perivascular areas. A,** Representative images showing GFAP staining (green) within the tumor core and periphery of *Ntv-a;Aqp4*$^{+/+}$ and *Ntv-a;Aqp4*$^{-/-}$ mice bearing Nf1 mGBM. DAPI was used as a nuclear stain. Images were taken on the Leica SP8 confocal microscope. Scale bar = 100 um**. B**, Quantification of GFAP-positive area within the tumor core (left) and periphery (right) of *Ntv-a;Aqp4*$^{+/+}$ and *Ntv-a;Aqp4*$^{-/-}$ mice bearing Nf1 mGBM. Two-tailed Student's t-test, *p<0.05. **C**, Schematic illustration of perivascular GFAP staining qauntification in *Ntv-a;Aqp4*$^{+/+}$ and *Ntv-a;Aqp4*$^{-/-}$ mice bearing PDGFB mGBM. ROIs were outlined around CD31-positive blood vessels within the tumor core for the purpose of quantification and kept consistent across samples. **D**, Representative images of CD31-positive blood vessels (red) and GFAP-positive perivascular astrocytes (green) within the tumor core of *Ntv-a;Aqp4*$^{+/+}$ and *Ntv-a;Aqp4*$^{-/-}$ mice bearing PDGFB mGBM. DAPI (blue) was used as a nuclear stain. Images were taken on the Leica SP8 confocal microscope. Scale bar=50 um. **E**, Quantification of perivascular GFAP-positive area in *Ntv-a;Aqp4*$^{+/+}$ and *Ntv-a;Aqp4*$^{-/-}$ mice bearing PDGFB mGBM. Two-tailed Student's t-test, **p<0.01.

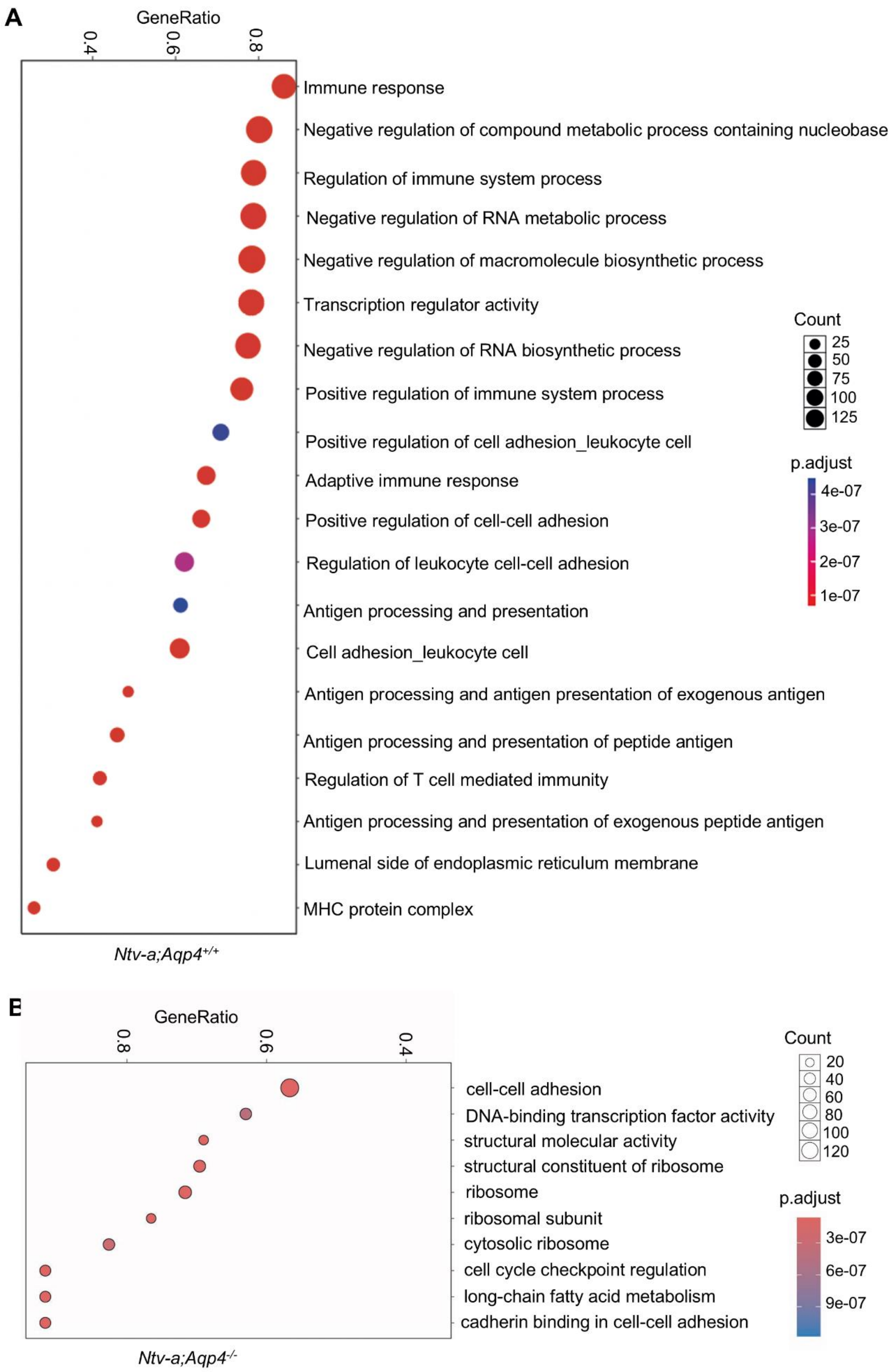
A
GeneRatio
0.4
0.6
0.8
Immune response
Negative regulation of compound metabolic process containing nucleobase
Regulation of immune system process
Negative regulation of RNA metabolic process
Negative regulation of macromolecule biosynthetic process
Transcription regulator activity
Negative regulation of RNA biosynthetic process
Positive regulation of immune system process
Positive regulation of cell adhesion_leukocyte cell
Adaptive immune response
Positive regulation of cell-cell adhesion
Regulation of leukocyte cell-cell adhesion
Antigen processing and presentation
Cell adhesion_leukocyte cell
Antigen processing and antigen presentation of exogenous antigen
Antigen processing and presentation of peptide antigen
Regulation of T cell mediated immunity
Antigen processing and presentation of exogenous peptide antigen
Lumenal side of endoplasmic reticulum membrane
MHC protein complex
Count
25
50
75
100
125
p.adjust
4e-07
3e-07
2e-07
1e-07
Ntv-a;Aqp4+/+
B
GeneRatio
0.8
0.6
0.4
cell-cell adhesion
DNA-binding transcription factor activity
structural molecular activity
structural constituent of ribosome
ribosome
ribosomal subunit
cytosolic ribosome
cell cycle checkpoint regulation
long-chain fatty acid metabolism
cadherin binding in cell-cell adhesion
Count
20
40
60
80
100
120
p.adjust
3e-07
6e-07
9e-07
Ntv-a;Aqp4-/-

109

**Supplementary Fig.6. Gene set enrichment analysis of differentially expressed genes in the astrocyte cluster from *Aqp4*-proficient and deficient tumor-bearing mice.** **A,** Dot plot showing gene set enrichment analysis of differentially expressed genes within the astrocyte cluster in *Ntv-a;Aqp4*$^{+/+}$ mice (n=4) bearing PDGFB mGBM. **B,** Dot plot showing gene set enrichment analysis of differentially expressed genes within the astrocyte cluster in *Ntv-a;Aqp4*$^{-/-}$ mice (n=3) bearing PDGFB mGBM.

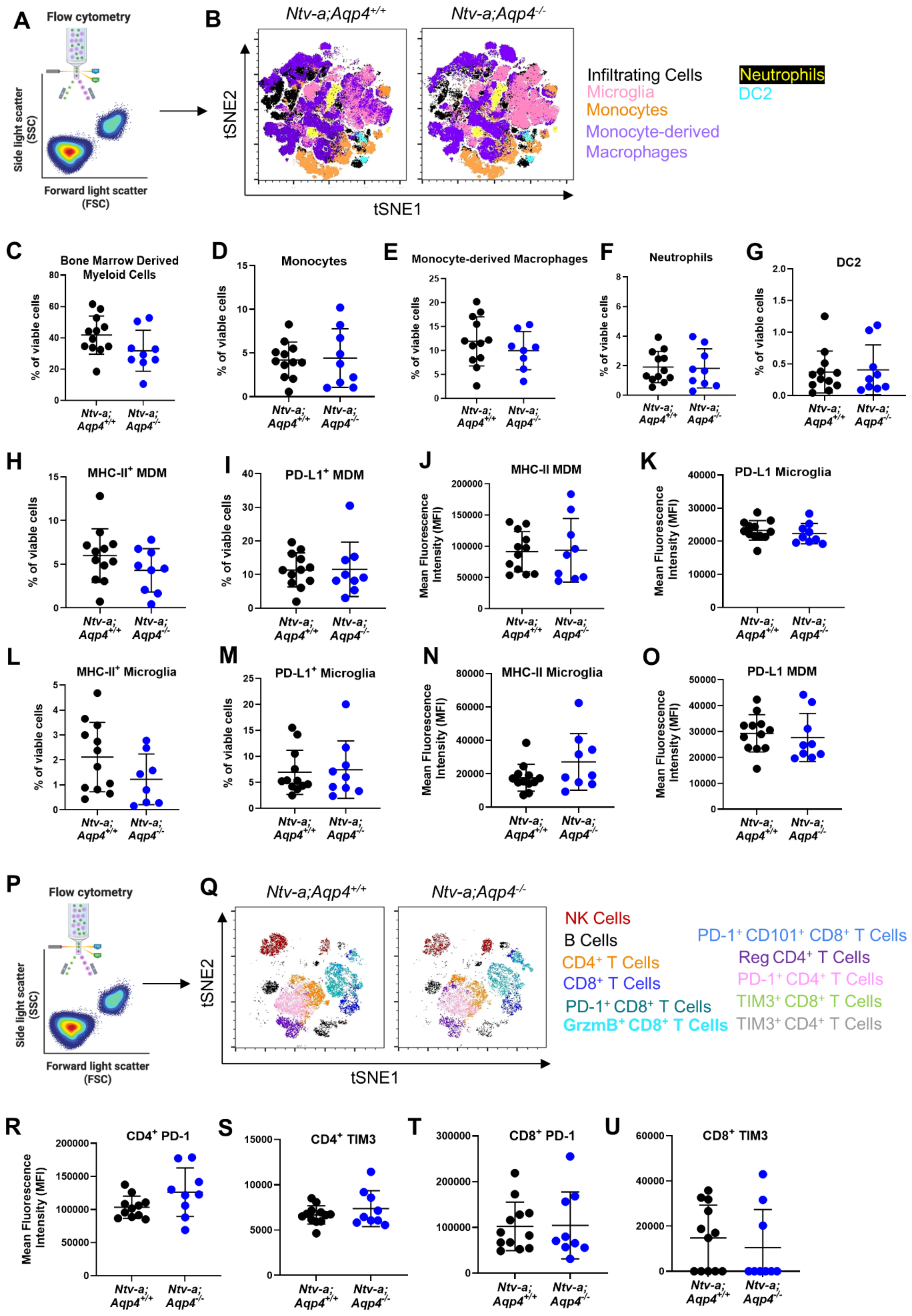
A
Flow cytometry
Side light scatter (SSC)
Forward light scatter (FSC)
B
Ntv-a;Aqp4+/+
Ntv-a;Aqp4-/-
tSNE2
tSNE1
Infiltrating Cells
Microglia
Monocytes
Monocyte-derived Macrophages
Neutrophils
DC2
C
Bone Marrow Derived Myeloid Cells
D
Monocytes
E
Monocyte-derived Macrophages
F
Neutrophils
G
DC2
% of viable cells
H
MHC-II+ MDM
I
PD-L1+ MDM
J
MHC-II MDM
K
PD-L1 Microglia
Mean Fluorescence Intensity (MFI)
L
MHC-II+ Microglia
M
PD-L1+ Microglia
N
MHC-II Microglia
O
PD-L1 MDM
P
Flow cytometry
Q
NK Cells
B Cells
CD4+ T Cells
CD8+ T Cells
PD-1+ CD8+ T Cells
GrzmB+ CD8+ T Cells
PD-1+ CD101+ CD8+ T Cells
Reg CD4+ T Cells
PD-1+ CD4+ T Cells
TIM3+ CD8+ T Cells
TIM3+ CD4+ T Cells
R
CD4+ PD-1
S
CD4+ TIM3
T
CD8+ PD-1
U
CD8+ TIM3
Ntv-a; Aqp4+/+
Ntv-a; Aqp4-/-

**Supplementary Fig.7. Flow cytometric analysis of various myeloid and lymphoid populations in *Aqp4*-proficient and deficient tumor-bearing mice. A and B,** Schematic illustration of flow cytometry followed by tSNE plot showing proportion of myeloid subpopulations in *Ntv-a;Aqp4*$^{+/+}$ and *Ntv-a;Aqp4*$^{-/-}$ mice bearing PDGFB mGBM. **C-G**, Graphs showing quantification of various myeloid subpopulations gated by flow cytometry as a percentage of viable cells. **H,I,L and M**, Graphs showing quantification of MHC-II+ and PD-L1+ monocyte-derived macrophages (MDM) and microglia gated by flow cytometry as a percentage of viable cells. **J,K,N and O,** Graphs showing quantification of the mean fluorescence intensity (MFI) of MHC-II and PD-L1 expression on MDMs and microglia. **P and Q**, tSNE plot showing distribution of various lymphoid subpopulations in *Ntv-a;Aqp4*$^{+/+}$ and *Ntv-a;Aqp4*$^{-/-}$ mice bearing PDGFB mGBM. **R-U**, Graphs showing mean fluorescence intensity (MFI) of phenotypic markers PD-1 and TIM3 in $CD4^+$ and $CD8^+$ T cell populations in *Ntv-a;Aqp4*$^{+/+}$ and *Ntv-a;Aqp4*$^{-/-}$ mice bearing PDGFB mGBM.

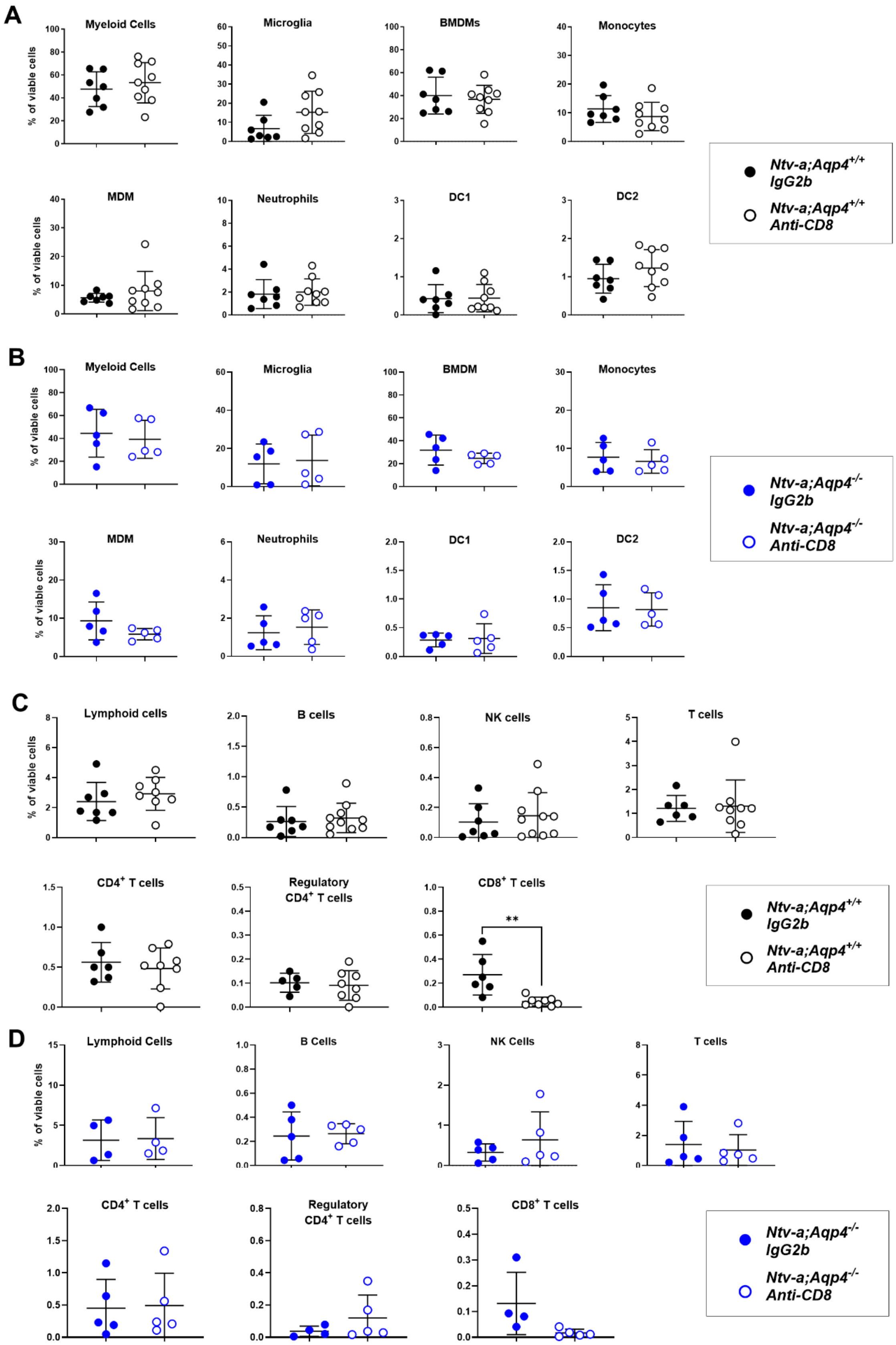
A
Myeloid Cells
Microglia
BMDMs
Monocytes
% of viable cells
MDM
Neutrophils
DC1
DC2
Ntv-a;Aqp4+/+ IgG2b
Ntv-a;Aqp4+/+ Anti-CD8
B
Myeloid Cells
Microglia
BMDM
Monocytes
MDM
Neutrophils
DC1
DC2
Ntv-a;Aqp4-/- IgG2b
Ntv-a;Aqp4-/- Anti-CD8
C
Lymphoid cells
B cells
NK cells
T cells
CD4+ T cells
Regulatory CD4+ T cells
CD8+ T cells
**
Ntv-a;Aqp4+/+ IgG2b
Ntv-a;Aqp4+/+ Anti-CD8
D
Lymphoid Cells
B Cells
NK Cells
T cells
CD4+ T cells
Regulatory CD4+ T cells
CD8+ T cells
Ntv-a;Aqp4-/- IgG2b
Ntv-a;Aqp4-/- Anti-CD8

**Supplementary Fig.8. Evaluation of the effect of CD8$^{+}$ T cell depletion on various immune populations in tumors from *Aqp4*-proficient and deficient mice. A,** Flow cytometry plots showing a comparison between various myeloid populations in *Ntv-a;Aqp4*$^{+/+}$ mice bearing PDGFB mGBM treated with IgG2b control or anti-CD8 antibody. **B**, Flow cytometry plots showing a comparison between various myeloid populations in *Ntv-a;Aqp4*$^{-/-}$ mice bearing PDGFB mGBM treated with IgG2b control or anti-CD8 antibody. **C,** Flow cytometry plots showing a comparison between various lymphoid populations in *Ntv-a;Aqp4*$^{+/+}$ mice bearing PDGFB mGBM treated with IgG2b control or anti-CD8 antibody. **D**, Flow cytometry plots showing a comparison between various lymphoid populations in *Ntv-a;Aqp4*$^{-/-}$ mice bearing PDGFB mGBM treated with IgG2b control or anti-CD8 antibody. All quantifications were compared using two-tailed Student's t-test.

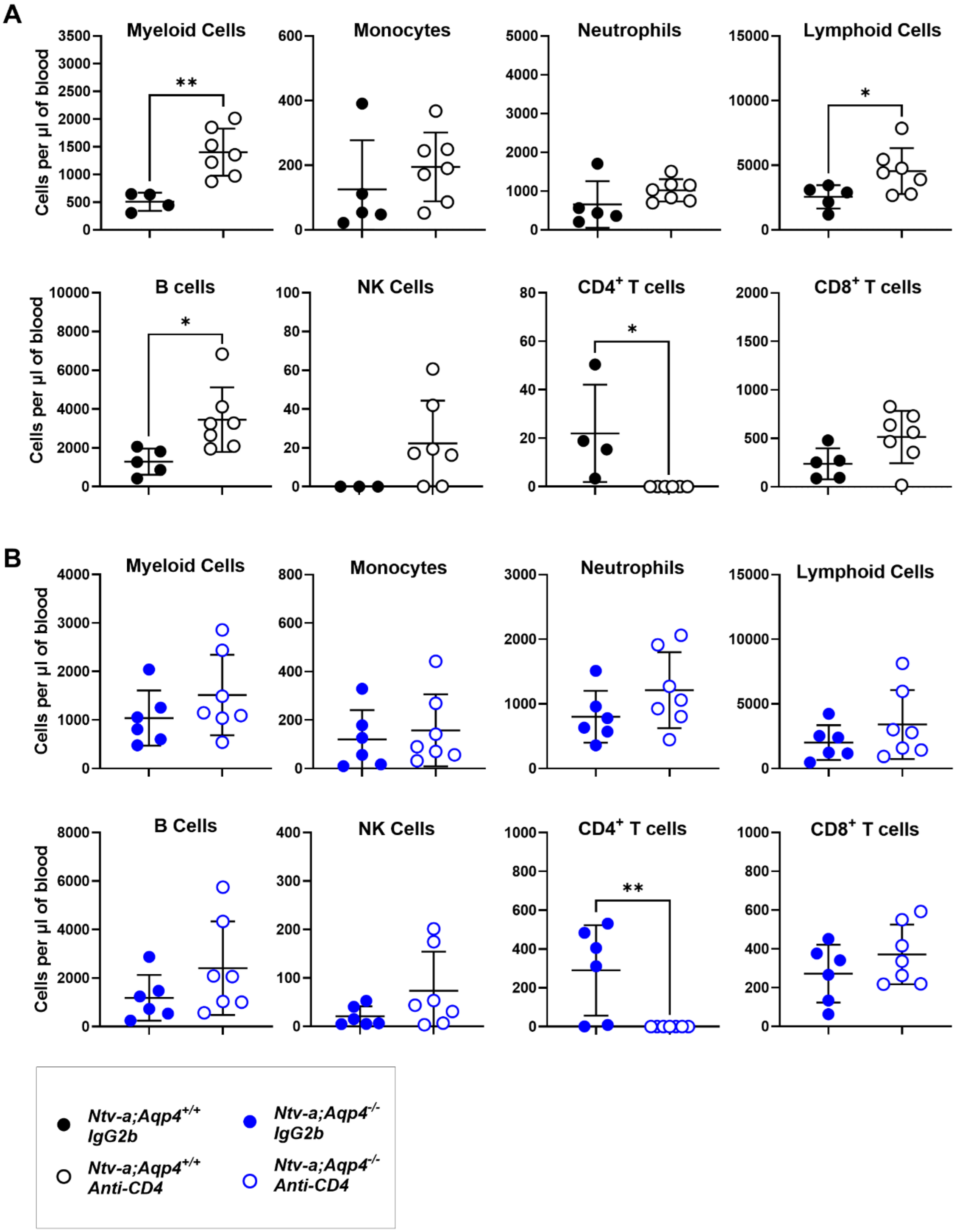


**Supplementary Fig.9. Evaluation of the effect of CD4$^+$ T cell depletion on various immune populations in the blood of *Aqp4*-proficient and deficient tumor-bearing mice. A,** Flow cytometry plots showing quantification of various immune cell populations in the blood of *Ntv-*

*a;Aqp4*$^{+/+}$ mice bearing PDGFB mGBM treated with IgG2b control or anti-CD4 antibody. **B**, Flow cytometry plots showing quantification of different immune cell populations in the blood of *Ntv-a;Aqp4*$^{-/-}$ mice bearing PDGFB mGBM treated with IgG2b control or anti-CD4 antibody. All quantifications were compared using two-tailed Student's t-test.

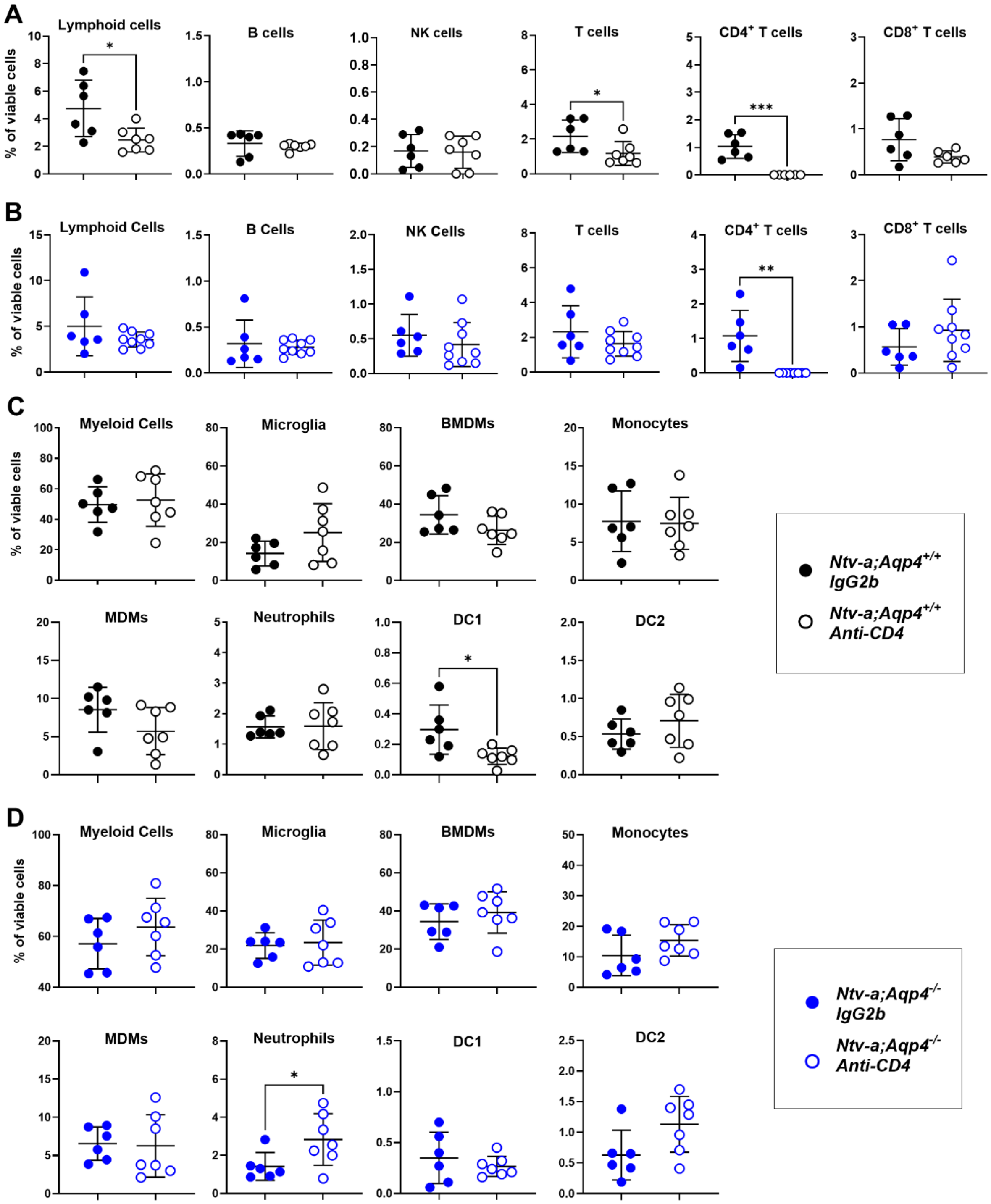


**Supplementary Fig.10. Evaluation of the effect of CD4$^{+}$ T cell depletion on various immune populations in tumors from *Aqp4*-proficient and deficient mice. A,** Flow cytometry plots showing a comparison between various lymphoid populations in *Ntv-a;Aqp4*$^{+/+}$ mice bearing

PDGFB mGBM treated with IgG2b control or anti-CD4 antibody. **B**, Flow cytometry plots showing a comparison between various lymphoid populations in *Ntv-a;Aqp4*$^{-/-}$ mice bearing PDGFB mGBM treated with IgG2b control or anti-CD4 antibody. **C**, Flow cytometry plots showing a comparison between various myeloid populations in *Ntv-a;Aqp4*$^{+/+}$ mice bearing PDGFB mGBM treated with IgG2b control or anti-CD4 antibody. **D**, Flow cytometry plots showing a comparison between various myeloid populations in *Ntv-a;Aqp4*$^{-/-}$ mice bearing PDGFB mGBM treated with IgG2b control or anti-CD4 antibody. All quantifications were compared using two-tailed Student's t-test.

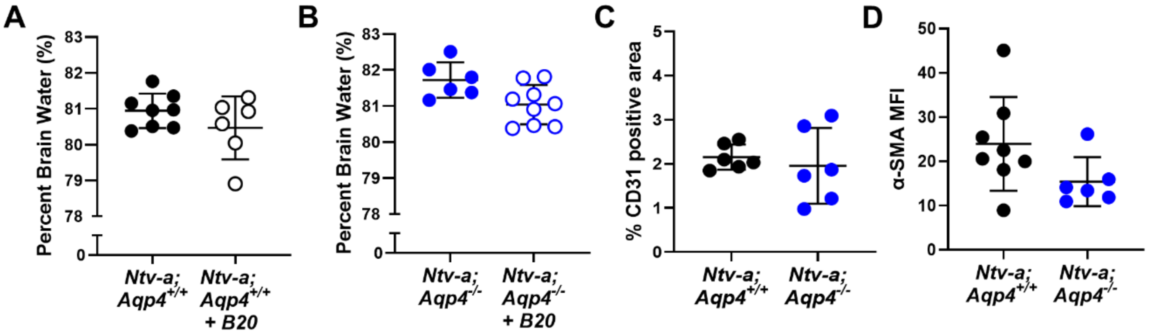


**Supplementary Fig. 11. Effects of anti-VEGF treatment on edema content and vasculature in *Aqp4*-proficient and deficient tumor-bearing mice. A,** Graph showing percent brain water calculated by a wet/dry assay in *Ntv-a;Aqp4*$^{+/+}$ mice bearing PDGFB mGBM, with B20-4.1.1 treatment (n=8) and 0.9% saline control treatment (n=6)**. B**, Graph showing percent brain water calculated by a wet/dry assay in *Ntv-a;Aqp4*$^{-/-}$ mice bearing PDGFB mGBM, with B20-4.1.1 treatment (n=6) and 0.9% saline control treatment (n=9)**. C**, Quantification of CD31 positive area (percentage) in *Ntv-a;Aqp4*$^{+/+}$ (n=6) and *Ntv-a;Aqp4*$^{-/-}$ (n=6) mice bearing PDGFB mGBM. **D**, Quantification of mean fluorescence intensity (MFI) for alpha-smooth muscle actin (alpha-SMA) in *Ntv-a;Aqp4*$^{+/+}$ (n=8) and *Ntv-a;Aqp4*$^{-/-}$ (n=6) mice bearing PDGFB mGBM. All quantifications were compared using two-tailed Student's t-test.

### Supplementary Tables

**Supplementary Table 1. Human GBM samples used for AQP4-edema correlation studies and Rarecyte ORION staining.**

| Sample ID | EGFR VIII | PDGFRA Amplification | NF1 | TERT |
|---|---|---|---|---|
| 21-0189 | Variant Not Detected | No Amplification | Mutated, Pathogenic | Mutation Present |
| 19-0051 | Variant Detected | No Amplification | Indeterminate | Indeterminate |
| 19-0142 | Variant Detected | No Amplification | Indeterminate | Indeterminate |
| 23-0170 | Variant Detected | No Amplification | No Mutation | Mutation Present |
| 22-0036 | Variant Detected | No Amplification | No Mutation | Indeterminate |
| 22-0169 | Variant Not Detected | No Amplification | Mutated, Pathogenic | Indeterminate |
| 23-0198 | Variant Detected | No Amplification, But Variant of Uncertain Significance | No Mutation | Mutation Present |
| 21-0057 | Variant Detected | No Amplification | No Mutation | Mutation Present |
| 23-0093 | Variant Detected | No Amplification | No Mutation | Mutation Present |
| 23-0336 | Variant Not Detected | Amplification Present | No Mutation | Mutation Present |

**Supplementary Table 2. Human GBM primary cultures and tissue samples used for Western Blots**

| Sample ID | Sample Type | Mutational profile |
|---|---|---|
| 24286 | Tumor tissue | IDH wt, EGFR amp positive (Sema4: EGFR 91 copies, TERT c-146C>T 43%, TP53 p.Y220D missense 66%); possible PTEN and RB1 loss; possible CDKN2A/B loss; |
| 29282 | Tumor tissue | IDH wt, EGFR amplification (55 copies); EGFR p.A289V; PTEN p.K332Tfs*8; TERT c.-124C>T |
| 27419 | Tumor tissue | IDH wt, NF1 splice site 2850+1G>A; CDKN2A loss; PTEN M199del; TERT -146C>T; TP53 T211_H214del; MS-Stable; 4 muts/ MB |
| 11849 | Tumor tissue | IDH wt, PDGFRA and KIT amplified |
| 31100 | Tumor tissue | IDH wt, CDKN2A loss; MTAP loss; NF1 N2387_F2388de; NF1 K1444E; PTEN loss; PTPN11 Y63C; TERT promoter -146C>T; MS-Stable; 1 mut/Mb |
| hAC1 | Adult frontal neocortex | Normal cortex |
| 827 | Glioma stem cell culture | Proneural subtype |
| 559 | Glioma stem cell culture | Mesenchymal subtype |
| 448 | Glioma stem cell culture | Proneural subtype |

**Supplementary Table 3. Murine GBM primary cultures and tissue samples used for Western Blots**

| Sample ID | Sample Type | Mutational profile |
|---|---|---|
| mPDGFB-1 | Tumor tissue | *PDGFB* overexpression, silenced *p53* |
| mPDGFB-2 | Tumor tissue | *PDGFB* overexpression, silenced *p53* |
| mPDGFB-3 | Tumor tissue | *PDGFB* overexpression, silenced *p53* |
| mNf1-1 | Tumor tissue | *PDGFA* overexpression, silenced *Nf1/p53/Pten* |
| mNf1-2 | Tumor tissue | *PDGFA* overexpression, silenced *Nf1/p53/Pten* |
| mNf1-3 | Tumor tissue | *PDGFA* overexpression, silenced *Nf1/p53/Pten* |
| mNaive-1 | Cortex Tissue | Normal cortex |
| mNaive-2 | Cortex Tissue | Normal cortex |
| 1° Astrocytes | Primary culture | Primary astrocyte cultures from naïve P0-P3 pups |
| mPDGFB-KO | Tumor tissue | *PDGFB* overexpression, silenced *p53*; *Aqp4* knockout |
| CT2A | Tumor cell line | *N-ras* mutation, *Cdkn2a/b* heterozygous deletion*, Pten* deficiency |
| GL261 | Tumor cell line | *K-Ras*/*Trp53* mutations, *c-myc* overexpression |
| 1861 | Primary culture | *Nf1/p53* silenced |
| 4622 | Primary Culture | *Nf1/p53* silenced |

**Supplementary Table 4. Immunohistochemistry and immunofluorescence antibodies used**

| Marker | Species | Dilution | Vendor | Cat# |
|---|---|---|---|---|
| GFAP | Mouse | 1:500 | Cell Signaling | 3670 |
| AQP4 | Rabbit | 1:100 | Millipore Sigma | AB3594 |
| GFP | Rabbit | 1:100 | ThermoFisher | A-11122 |
| CD31 | Rat | 1:50 | Dianova | DIA-310 |
| α-SMA | Rat | 1:500 | Dako | M0851 |
| pH3 | Rabbit | 1:200 | Sigma | 06-570 |

**Supplementary Table 5. Antibodies used for flow cytometry.**

**Myeloid panel (IC - intracellular)**

| Marker | Fluorophore | Dilution | Vendor | Cat# |
|---|---|---|---|---|
| CD45 | FITC | 1:100 | Invitrogen | 11-0451-81 |
| CD11b | PerCP-Cy5.5 | 1:100 | BD | 550993 |
| CD11c | APC | 1:100 | BD | 550261 |
| MHCII | A700 | 1:100 | Invitrogen | 56-5321-80 |
| CD103 | BUV395 | 1:100 | BD | 748253 |
| CD24 | BUV496 | 1:200 | BD | 612953 |
| PDL1 | BV605 | 1:100 | BD | 745135 |
| CX3CR1 | BV650 | 1:100 | Biolegend | 149033 |
| F4/80 | BV711 | 1:100 | Biolegend | 123147 |
| Ly6g | V450 | 1:100 | BD | 560603 |
| CXCR2 | PE | 1:100 | Biolegend | 149609 |
| CD49d | PE/D | 1:100 | Biolegend | 103625 |
| Ly6c | PE-Cy7 | 1:200 | BD | 560593 |
| Ki67 (IC) | eF506 | 1:50 | Invitrogen | 69-5698-80 |

**Lymphoid panel (IC - intracellular)**

| Marker | Fluorophore | Dilution | Vendor | Cat# |
|---|---|---|---|---|
| CD11b | PerCP-Cy5.5 | 1:100 | BD | 550993 |
| CD101 | APC | 1:100 | Invitrogen | 17-1011-80 |
| CD4 | APC-Cy7 | 1:100 | Biolegend | 100526 |
| B220 | BV605 | 1:100 | Biolegend | 103243 |
| NK1.1 | BV711 | 1:100 | BD | 740663 |
| PD1 | BV785 | 1:100 | Biolegend | 135225 |
| CD45 | BV510 | 1:100 | Biolegend | 103137 |
| CD8 | PE-Cy5 | 1:100 | BD | 553034 |

| | | | | |
|---|---|---|---|---|
| CD3 | PE/D | 1:100 | Biolegend | 100348 |
| Tim-3 | PE-Cy7 | 1:100 | Invitrogen | 25-5870-82 |
| GrzmB (IC) | PE | 1:50 | Invitrogen | 12-8899-41 |
| FoxP3 (IC) | FITC | 1:50 | Invitrogen | 11-5773-82 |
| Ki67 (IC) | eF506 | 1:50 | Invitrogen | 69-5698-80 |

**Blood panel**

| Marker | Fluorophore | Dilution | Brand | Cat# |
|---|---|---|---|---|
| CD45 | FITC | 1:100 | Invitrogen | 11-0451-81 |
| CD11b | PerCP-Cy55 | 1:100 | BD | 550993 |
| Ly6g | V450 | 1:100 | BD | 560603 |
| Ly6c | PE-Cy7 | 1:200 | BD | 560593 |
| CD4 | APC-Cy7 | 1:100 | Biolegend | 100526 |
| B220 | BV605 | 1:100 | Biolegend | 103243 |
| NK1.1 | BV711 | 1:100 | BD | 740663 |
| CD8 | BV510 | 1:100 | Biolegend | 100752 |
| CD3 | PE/D | 1:100 | Biolegend | 100348 |

**Supplementary Table 6. Orion multiplex IF antibody panel**

| Marker | ArgoFluor Dye | Species | Clone / Type | Cat# | Vendor |
|---|---|---|---|---|---|
| SPP1 (Osteopontin) | Argo520 | Goat | Polyclonal | 62-1022-501 | RareCyte |
| IBA1 | Argo548 | Rabbit | Polyclonal | 019-19741 | FUJIFILM Wako |
| GFAP | Argo660L | Mouse | 2.2B10 | 62-1251-601 | RareCyte |
| CD163 | Argo602 | Rabbit | EPR14643 | 62-1006-601 | RareCyte |
| CD8α | Argo624 | Mouse | AMC908 | 62-1007-601 | RareCyte |
| OLIG2 | Argo662 | Rabbit | EPR2673 | ab220796 | Abcam |
| AQP4 | Argo676 | Rabbit | Polyclonal | 16473-1-PBS | Proteintech |
| CD4 | Argo706 | Rabbit | EPR6855 | 62-1098-701 | RareCyte |
| PD-L1 | Argo724 | Mouse | 73-10 | 62-1066-701 | RareCyte |
| CD15 | Argo760 | Mouse | QA19A61 | 62-1048-701 | RareCyte |
| CD20 | Argo782 | Mouse | L26 | 62-1240-601 | RareCyte |
| CD31 | Argo812 | Rabbit | EPR3094 | 62-1045-701 | RareCyte |
| ALDH1L1 | Argo845 | Rabbit | EPR12744(B) | ab249853 | Abcam |
